# Autism-specific PTEN p.I135L mutation and an autism genetic background combine to dysregulate cortical neurogenesis

**DOI:** 10.1101/2022.11.15.516317

**Authors:** Shuai Fu, Luke Bury, Jaejin Eum, Anthony Wynshaw-Boris

**Affiliations:** Department of Genetics and Genome Sciences, Case Western Reserve University, Cleveland, OH 44106; Department of Molecular Medicine, Cleveland Clinic Lerner College of Medicine, Cleveland, OH, 44195

**Keywords:** PTEN, ASD, CRISPR-Cas9 genome editing, iPSC, Cortical organoids

## Abstract

Alterations in cortical neurogenesis are implicated in neurodevelopmental disorders including autism spectrum disorders (ASDs). The contribution of genetic backgrounds, in additional to ASD risk genes, on cortical neurogenesis remain understudied. Here, using isogenic induced pluripotent stem cell (iPSC)-derived neural progenitor cells (NPCs) and cortical organoid models, we report that a heterozygous *PTEN* p.I135L mutation found in an ASD patient with macrocephaly activates PI3K/AKT and dysregulates cortical neurogenesis in an ASD genetic background-dependent fashion. Transcriptome analysis at both bulk and single cell level revealed *PTEN* p.I135L mutation and ASD genetic background affected genes involved in neurogenesis, neural development and synapse signaling. We also found that this *PTEN* p.I135L mutation led to overproduction of NPC subtypes as well as neuronal subtypes including both deep and upper layer neurons in its ASD background, but not when introduced into a control genetic background. These findings provide experimental evidence that both a *PTEN* p.I135L mutation and ASD genetic background contribute to cellular features consistent with ASD associated with macrocephaly.

## INTRODUCTION

ASDs are a group of phenotypically complex and genetically heterogeneous neurodevelopmental disorders^1^. Mutations in nearly 1000 genes have been associated with ASD^2,3^. Many of these gene mutations are de novo, but a role for inherited common variation is likely to be a major contributor to ASD genetic risk^4^. Approximately 20% of ASD individuals display early brain overgrowth^5,6^ and 15% of ASD subjects with early brain overgrowth possess mutations in the gene *PTEN*^7^. *PTEN* is a well-known tumor suppressor gene which also acts as a lipid phosphatase that dephosphorylates PIP3 to PIP2, reducing PI3K/AKT activation, and leading to reduced cell proliferation or increased apoptosis^8,9^. Compared to tumor-related *PTEN* mutant forms that result in cancer prone familial risk, a majority of ASD-associated *PTEN* mutations do not substantially disrupt the lipid phosphatase function of *PTEN*^10^. In addition, subjects with *PTEN* mutations do not invariably display ASD, and many ASD subjects do not carry mutations in known ASD risk genes. This provides further support for the hypothesis that, in addition to key ASD risk genes such as *PTEN*, ASD patient genetic background—the spectrum of variants and mutations found in throughout the genome and/or epigenome—may also contribute to ASD pathology^11,12^. However, experimental support for this hypothesis is lacking.

Here, we have utilized iPSC models and genome editing to address this question. iPSCs have been extensively used for neurodevelopmental disease modeling^13–17^. iPSCs can be used to produce NPCs and neurons in two-dimensional culture, as well as three-dimensional self-organizing brain organoids^18^, which mimic the neurogenesis trajectory during human fetal brain development^19,20^. Bi-directional genome editing using CRISPR-Cas9 on iPSCs derived from both healthy control and ASD subjects enables the effects of key ASD risk genes as well as the contribution of the genetic background to ASD etiology to be determined.

We previously reported increased NPC proliferation in eight iPSC-derived NPCs from patients with early brain overgrowth, compared with five control lines^16^. We identified a heterozygous *PTEN* p.I135L mutation among other variants in one of the eight ASD subjects through whole exome sequencing^16^. Therefore, we produced isogenic lines with wild-type *PTEN* (*PTEN* WT/WT), the ASD *PTEN* mutation (*PTEN* WT/I135L), and complete disruption of *PTEN* (*PTEN* KO/KO) in both ASD and control backgrounds. Using NPC cultures and self-organizing organoids, we found that the *PTEN* p.I135L ASD heterozygous mutation led to increased NPC proliferation, enlarged organoid size, dysregulated genes related to neurogenesis, and, specifically on the ASD genetic background, the overproduction of deep and upper layer neurons. These effects were modified by the genetic background, since the ASD genetic background itself also led to increased NPC proliferation and dysregulation of genes related with neurogenesis depending on the *PTEN* genotype. Furthermore, we observed *PTEN* WT/I135L organoids displayed accelerated upper layer neuronal maturation in the autistic background. Lastly, we identified that both *PTEN* p.I135L and ASD genetic background led to dysregulated synaptic signaling in multiple neuronal subtypes. These studies provide strong experimental evidence that both a specific ASD mutation and ASD genetic background are important for cellular surrogates of brain overgrowth in ASD.

## RESULTS

### *PTEN* p.I135L mutation leads to increased NPC proliferation in both control and ASD genetic background

To test the effect of the ASD *PTEN* p.I135L mutation and genetic background on the cellular phenotypes associated with ASD, we generated isogenic *PTEN* iPSCs in both ASD and control backgrounds. We corrected the *PTEN* p.I135L patient mutation in the ASD iPSC line (referred to as “Apex”) containing this mutation using CRISPR-Cas9^D10A^ double nickase genome editing^21^. We also introduced the same heterozygous *PTEN* p.I135L mutation into the control iPSC line (referred to as “Chap”, Figure S1A, B). In addition, we abolished *PTEN* expression by creating *PTEN* KO/KO lines in both control (Chap) and ASD (Apex) genetic backgrounds (Figure 1B). We validated our CRISPR-edited clones using Sanger (Figure S1C) and Miseq (Figure S1D) amplicon sequencing. We confirmed that our edited iPSCs were isogenic with the parental lines based on Short Tandem Repeat (STR) profiles (data not shown) and were karyotypically normal using either Nano-string human karyotype panel or traditional G-banding (Figure S1E, F). Thus, we produced two sets of isogenic PTEN iPSC lines: one set on the control background (Chap *PTEN* WT/WT, Chap *PTEN* WT/I135L and Chap *PTEN* KO/KO) and one set on the ASD genetic background (Apex *PTEN* WT/WT, Apex *PTEN* WT/I135L and Apex *PTEN* KO/KO).

**Figure 1.**
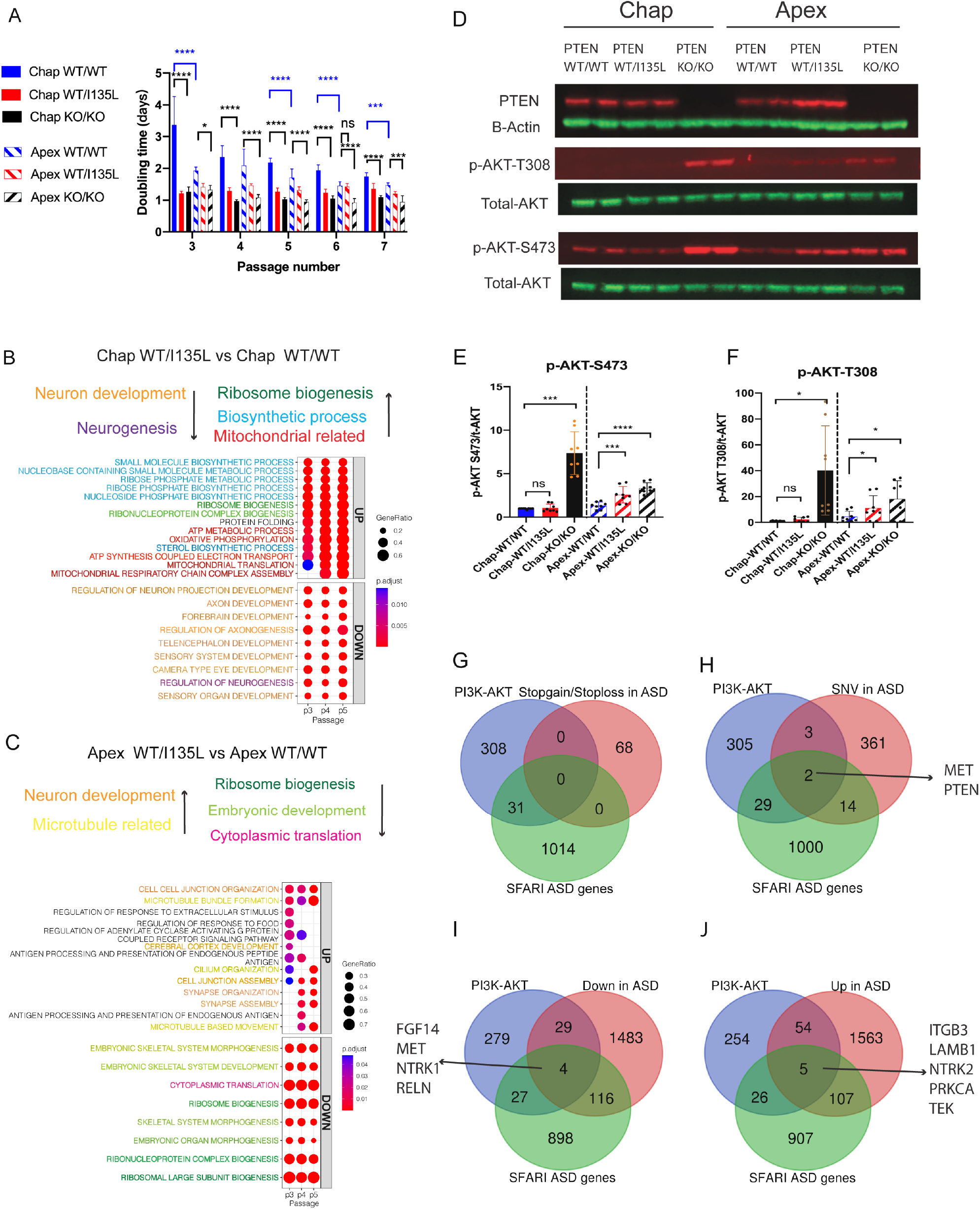
*PTEN* WT/I135L mutation dysregulates genes important for neurogenesis and impairs canonical *PTEN* activity in the ASD but not in the control genetic background. (A) Isogenic *PTEN* panel iPSCs were differentiated to NPCs. From passages 2 to 6, cells were plated at the same density and population doubling time at each passage was calculated. Results of all lines (3 independent cell culture replicates per line) are presented as mean±sd. P values were calculated using one-way anova with Sidak’s multiple comparisons. * p <0.05, ** p<0.01, *** p<0.001, **** p<0.0001, ns, not significant. (B) GSEA analysis for the effect of *PTEN* WT/I135L in the control background. The comparecluster function was used to run GSEA for multiple passages at the same time, with showCategory = 5. P values were adjusted using the Benjamini-Hochberg correction. (C) GSEA analysis for the effect of *PTEN* WT/I135L in the ASD genetic background. The comparecluster function was used to run GSEA for multiple passages at the same time, with showCategory = 6. P values were adjusted using the Benjamini-Hochberg correction.(D) Representative western blot results, 10μg lysates from isogenic *PTEN* NPCs at passage 5 were used for immunoblot for *PTEN*, beta-actin and p-AKT Ser473, p-AKT-Thr308. (E) Quantification of the ratio of p-AKT S473 to total AKT, n=8, 4 independent experiments were performed and each with duplicate sample loading. (F) Quantification of the ratio of p-AKT T308 to total AKT, n=8, 4 independent experiments, duplicate samples were loaded for each experiment. For both (E) and (F), Results are presented as mean±sd, P values were calculated using repeated measure one-way anova with Sidak’s multiple comparisons. * p <0.05, ** p<0.01, *** p<0.001, **** p<0.0001, ns, not significant. (G) Venn diagram for the identification of the overlapping genes among 339 PI3K/AKT pathway gene sets, 1045 SFARI ASD genes and 68 stop gains or stop loss mutations in Apex. (H) Venn diagram for the identification of the overlapping genes among 339 PI3K/AKT pathway gene sets, 1045 SFARI ASD genes and 380 SNVs in Apex. (I) Venn diagram for the identification of the overlapping genes among 339 PI3K/AKT pathway gene sets, 1045 SFARI ASD genes and 1632 downregulated genes in Apex. (J) Venn diagram for the identification of the overlapping genes among 339 PI3K/AKT pathway gene sets, 1045 SFARI ASD genes and 1729 upregulated genes in Apex.

To first test the effect of the ASD *PTEN* p.I135L mutation and genetic background on the NPC proliferation phenotype, we differentiated the three control and three ASD isogenic lines into dorsal forebrain NPCs expressing the markers PAX6 and FOXG1 (Figure S2). We measured proliferation using the population doubling time assay. The *PTEN* WT/I135L NPCs on both control and ASD genetic backgrounds displayed increased NPC proliferation as indicated by the significantly decreased doubling time across multiple passages (Figure 1A). Similar effects on NPC proliferation were observed when we abolished the expression of *PTEN* in both control and ASD genetic background, suggesting that the point mutation provides sufficient reduction of PTEN activity compared to the complete PTEN knock-out for accelerated NPC proliferation. Additionally, the Apex *PTEN* WT/WT NPCs proliferated faster than Chap PTEN WT/WT NPC (Figure 1A), indicating that ASD genetic background was also important for increased NPC proliferation (Figure 1A).

### *PTEN* p.I135L mutation dysregulates neurogenesis in NPCs as assessed by RNA-seq

To examine the effects of *PTEN* p.I135L mutation at the transcriptomic level, we then performed bulk RNA-sequencing (RNA-seq) on NPCs derived from each of the isogenic *PTEN* iPSC lines. In total, 54 RNA samples were sequenced: three experimental replicates for each isogenic iPSC-derived NPC group across 3 different passages averaging 30M paired-end reads per sample. We employed Gene Set Enrichment Analysis (GSEA) to provide sensitive and unbiased gene expression profiles for each *PTEN* genotype and genetic background^22,23^. We then focused on the top enriched Gene Ontology (GO) terms due to *PTEN* mutations in each genetic background across all passages for analysis. The *PTEN* WT/I135L NPCs in the control genetic background led to downregulated genes enriched for GO terms related with **neurogenesis** (regulation of neurogenesis) and **neuron development** (regulation of neuron projection development, axon development, forebrain development, regulation of axonogenesis, telencephalon development, sensory system development, camera type eye development and sensory organ development). We identified upregulated genes enriched for GO terms related with **mitochondria related** processes (ATP metabolic process, oxidative phosphorylation, ATP synthesis coupled electron transport, mitochondrial translation and mitochondrial respiratory chain complex assembly) and upregulated genes enriched for GO terms related with **biosynthetic process** (e.g., nucleoside phosphate biosynthetic process, sterol biosynthetic process, etc.) as well as GO terms related with **ribosome biogenesis** and **protein folding** (Figure 1B).

We performed similar GSEA analysis in *PTEN* WT/I135L NPCs in the ASD genetic background. Downregulated genes enriched for GO terms related to **embryonic development** (embryonic organoid morphogenesis, and embryonic skeletal system development) were found. In addition, we identified downregulated genes enriched for GO terms related with **cytoplasmic translation**, and upregulated genes enriched for GO terms related with **microtubules**. Interestingly, upregulated genes in the control genetic background were enriched for GO terms related with **ribosome biogenesis**, while in the ASD genetic background, genes enriched for the same GO terms were downregulated in the ASD genetic background due to *PTEN* WT/I135L mutation. Strikingly, GO terms related to **neuron development** (cerebral cortex development) which were enriched by downregulated genes in the control background were enriched by upregulated genes in the ASD genetic background (Figure 1B, C). These results suggest that gene ontology enrichment due to the *PTEN* WT/I135L mutation is dependent on the genetic background, and the ASD genetic background can even reverse the direction of GO terms enriched due to this *PTEN* WT/I135L mutation in the control background.

### *PTEN* p.I135L mutation activates PI3K/AKT in the ASD genetic background but not control background in 2D NPC culture

To test whether the p.I135L mutation impairs *PTEN* function, we performed western blot using lysates from NPC passage 5, passage 6 and passage 7 of all six isogenic *PTEN* NPCs, and looked for read outs of PTEN activity using anti-p-AKT Ser473 and Thr308 antibodies (Figure 1D-F, Figure S3A, B). As expected, *PTEN* KO/KO led to the complete loss of PTEN and dramatically decreased PTEN activity, indicated by increased p-AKT-T308 and p-AKT-S473 levels in both control and ASD genetic backgrounds. The Apex *PTEN* WT/I135L NPCs in the ASD background displayed a slight decrease of PTEN canonical activity indicated by very mild increases in p-AKT-T308 and p-AKT-S473 levels compared to *PTEN* KO/KO in the Apex ASD genetic background (Figure 1D-F). To our surprise, we found that Chap *PTEN* WT/I135L NPCs in the control background displayed no effect on the levels of both p-AKT-S473 and p-AKT-T308 (Figure 1D-F), suggesting that the *PTEN* WT/I135L mutation affects PTEN activity differently in control versus ASD genetic backgrounds.

### Genetic variants and differentially expressed genes in the ASD genetic background

To gain insight into the ASD background-dependent effect of the *PTEN* p.I135L point mutation on the PTEN activity as well as the differential gene expression findings (Figure 1), we first examined the whole exome sequencing (WES) data for the ASD line Apex^16^. There were 68 genes with either stop-loss or stop-gain mutations in Apex, and none of these overlapped with the 339 genes in the PI3K-AKT pathway derived from Molecular Signature Database (MsigDB) or the 1045 ASD Simons Foundation Autism Research Initiative (SFARI) risk genes (Figure 1G). In addition, 380 genes with SNVs were identified in WES data from Apex. Of these, two overlapped with both PI3K-AKT pathway and SFARI risk genes: *PTEN*, as expected, and *MET* (Figure 1H). An additional 3 SNVs overlapped with PI3K-AKT pathway genes while 14 genes overlapped with 1045 SFARI risk genes but were not in the PI3K-AKT pathway. Next, we determined whether any of the differentially expressed genes (DEGs), identified comparing autism vs control, are both part of the PI3K-AKT pathway, and also SFARI autism risk genes. Out of 1632 genes downregulated Apex vs Chap (with p(adjust) <0.05, and log2 fold change >=0.6), 33 genes belong to PI3K-AKT pathway, of which 4 are also SFARI ASD risk genes: *FGF14, MET, NTRK1* and *RELN* (Figure 1I). All of these genes are outstanding candidates that alone or together modulate the activity of PTEN in the ASD background. *FGF14* is a gene that expresses a brain-specific Fibroblast Growth Factor, and mutations in this gene are responsible for an autosomal dominant form of cerebellar ataxia^24^. *MET* is a tyrosine kinase important for stem cell growth and epithelial-mesenchymal interactions^25^. Besides being downregulated in the ASD background, it harbors an SNV. *NTRK1* is a neurotropin (NGF) receptor that is critical for neuronal development. Mutations in this gene result in cognitive disability^26^. *RELN* is a ligand for a number of receptors during development and adulthood, while mutations result in defects in neuronal migration and neurogenesis^27^. An additional 116 downregulated genes are also SFARI ASD risk genes. Among 1729 genes that are upregulated Apex vs Chap, 59 genes belonged to PI3K-AKT pathway, of which 5 were SFARI ASD risk genes: *ITGB3, LAMB1, NTRK2, PRKCA* and *TEK* (Figure 1J). Likewise, these genes are outstanding candidates that alone or together modulate the activity of PTEN in the ASD background. *ITGB3* codes for integrin beta-3, a cell adhesion molecule that has important synaptic functions during development^28^. *LAMB1* codes for laminin beta-1, and mutations in this gene cause cobblestone lissencephaly^29^. *NTRK2* is a neurotropin (NT-3) receptor that is critical for neuronal development. Mutations in this gene result in obesity^30^ and mood disorders^31^. *PRKCA* codes for protein kinase C-alpha^32^ that has been associated with ASD^33^, while *TEK* codes for TIE2, and is expressed almost exclusively in the vascular system^34^. In summary, these results suggest that ASD background contains dysregulated genes related to the PI3K/AKT pathway, providing an explanation for the ASD-background dependent effect of *PTEN* p.I135L mutation on PTEN activity.

### ASD genetic background dysregulates neurogenesis in NPCs as assessed by RNA-seq

Our experimental design also allowed us to examine the effects of ASD vs control genetic background with *PTEN* WT/WT, *PTEN* WT/I135L as well as *PTEN* KO/KO genotypes, focusing on the GSEA significant terms identified in an unbiased fashion (shown in Figure 1). For the *PTEN* WT/WT genotype, the ASD genetic background resulted in an enrichment of downregulated genes associated with GO terms related to **neurogenesis** (regulation of neurogenesis) and **neuron development** (cerebral cortex development, forebrain development and axon development) as well as microtubule related terms (Figure 2A). We also observed upregulated genes enriched for GO terms related with biosynthetic process, mitochondria translation, ribosome biogenesis, embryonic development, cytoplasmic translation and protein folding (Figure 2A).

**Figure 2.**
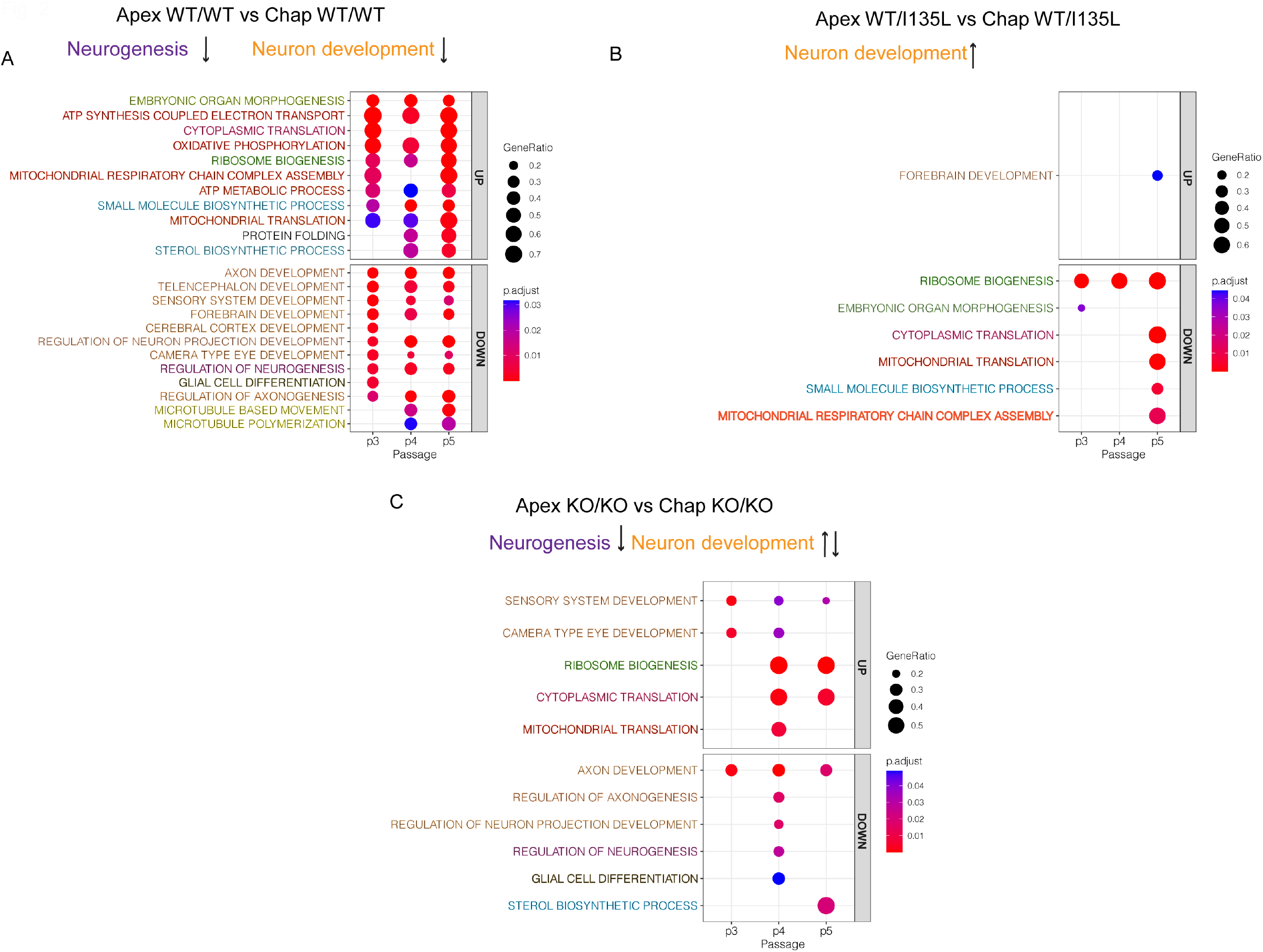
ASD genetic background dysregulates genes important for neurogenesis in 2D NPC culture. GSEA for the effect of ASD genetic background with *PTEN* WT/WT genotype (A), with *PTEN* WT/I135L genotype (B) and with *PTEN* KO/KO genotype (C). GO terms were preselected for visualization and p values were adjusted using the Benjamini-Hochberg correction.

For the *PTEN* WT/I135L genotype, the ASD genetic background affected similar GO terms, but the direction of the gene expression profile was often inverted. For example, genes enriched for GO terms related to “forebrain development” were now upregulated, while genes enriched for GO terms related to “ribosome biogenesis”, “mitochondria translation”, “biosynthetic process”, “embryonic organ morphogenesis” and “cytoplasmic translation” were now downregulated (Figure 2B). Furthermore, for the *PTEN* KO/KO genotype, the ASD genetic background affected genes enriched for GO terms related to **neuron development** in both positive (sensory system development) and negative (axon development, regulation of neuron projection development) directions (Figure 2C). These results suggest that the ASD genetic background affected similar pathways as the *PTEN* p.I135L mutation. Of note, ASD genetic background itself contributed to dysregulation of genes important for neurogenesis, in addition to the effects of the *PTEN* p.I135L mutation. Finally, as the severity of *PTEN* gene disruption increased from point mutation to complete loss-of-function, there were fewer GO terms altered in neurogenesis/neural development pathways selected in an unbiased fashion (Figure 1), suggesting that the severity of the *PTEN* mutation and the ASD background together contributed to the dysregulation of neural development.

### *PTEN* p.I135L mutation in ASD genetic background leads to overproduction of neural progenitors, deep and upper layer neurons in cortical organoids

To further interrogate the effect of *PTEN* p.I135L during cortical neurogenesis, we derived cortical organoids from our isogenic *PTEN* panel iPSCs using a dual SMAD and WNT inhibition protocol^14,35,36^, and cultured the organoids until week 21. We quantified organoid sizes and found that *PTEN* WT/I135L organoids displayed enlarged size in both control and ASD genetic background at week 4 (Figure 3A). However, at week 8, *PTEN* WT/I135L organoids continued enlarging in the control genetic background, while similar organoid enlargement was not observed in ASD background (Figure 3B). This suggests that the effect of this ASD-specific *PTEN* mutation on organoid growth was affected by ASD genetic background, consistent with findings in NPCs. In addition, organoids produced in the ASD genetic background were always larger than those from the control background regardless of the *PTEN* genotype. We fixed organoids and performed immunohistochemistry for NPC subtypes, as well as deep and upper layer neurons on the isogenic *PTEN* organoids between weeks 4 and 21 (Figure 3C, J, Figure S4B, S5B, S5D) (Figure S4A, S5A, S5C, S4C, S4D). As deep layer neurons and upper layer neurons can be generated from both intermediate progenitor cells (IPCs) and outer radial glia cells (oRGs)^37^, we then stained for TBR2, an IPC marker, and HOPX, an oRG-specific marker. Increased production of IPCs at weeks 4 and 10, as well as oRGs at week 10 (Figure 3D, E, K) were found in ASD *PTEN* WT/I135L vs. ASD *PTEN* WT/WT ASD background organoids. Surprisingly, no significant differences were observed for the *PTEN* WT/I135L mutation on IPCs or oRGs in the control genetic background. As expected from the IPC and oRG findings, we identified a significantly increased proportion of cells expressing deep layer neuron markers such as CTIP2 and TBR1 (Figure 3F,G, H, I), as well as the overproduction of upper layer neurons as indicated by SATB2 staining (Figure 3L, M). This was found in ASD *PTEN* WT/I135L vs ASD *PTEN* WT/WT ASD background organoids, but not in the Control *PTEN* WT/I135L vs Control *PTEN* WT/WT control background organoids. These results suggest that *PTEN* WT/I135L leads to increased production of NPC subtypes including IPCs and oRGs only in the ASD genetic background, leading to the overproduction of the deep layer and superficial layer neurons.

**Figure 3.**
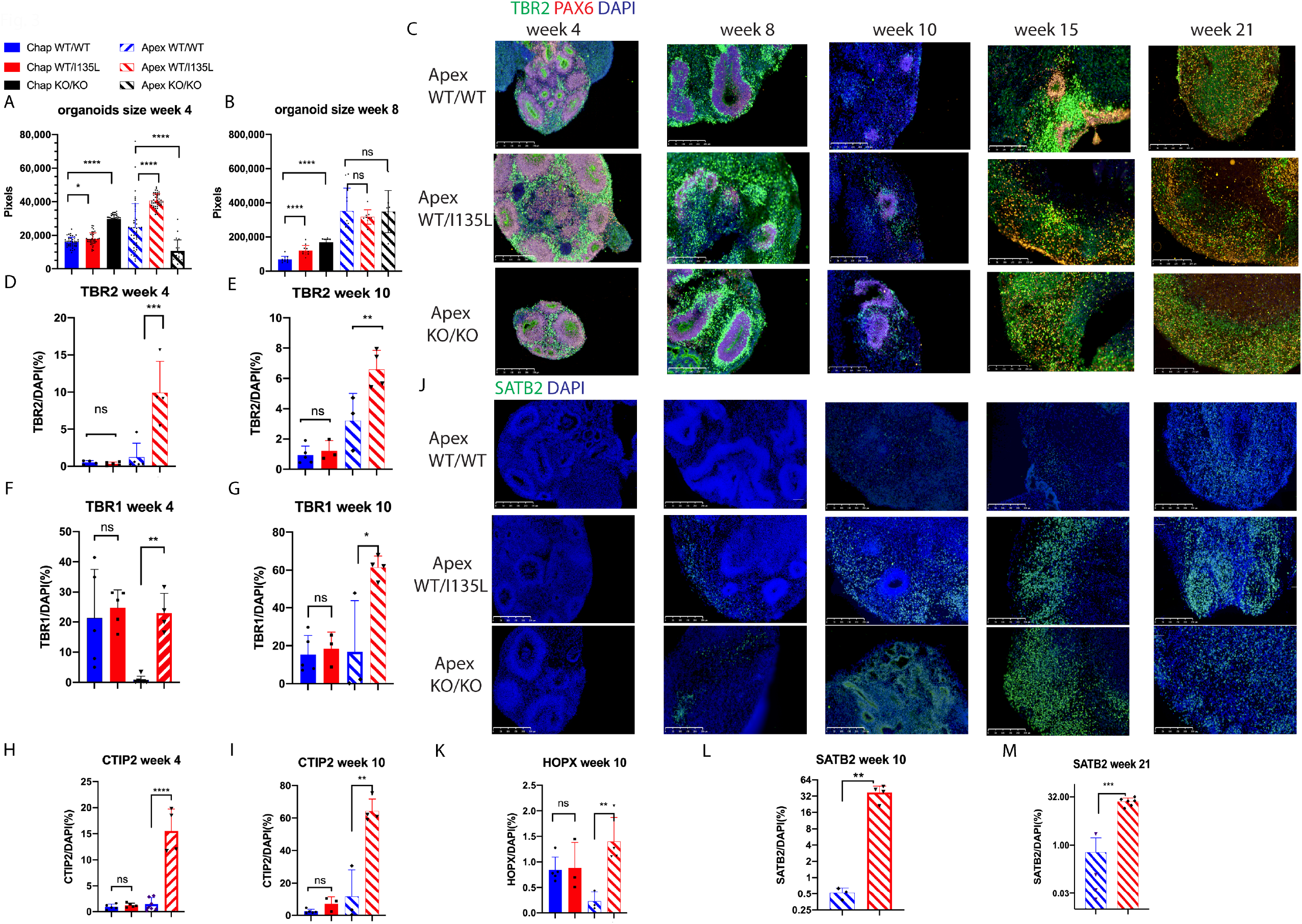
*PTEN* WT/I135L mutation leads to NPC subtypes and neuron overproduction in the ASD genetic background but not in the control genetic background. Organoid size quantification for week 4 (A) and week 8 (B). One-way anova was used, and p values were adjusted using Sidak’s multiple correction. * p<0.05, **** p<0.0001, ns, not significant. Representative IHC images for the isogenic *PTEN* organoids for week 4, week 8, week 10, week 15 and week 21 for the NPC marker PAX6, the IPC marker TBR2 (C), and the upper layer neuron marker SATB2 (j) scale bar 250μm. Quantification of TBR2/DAPI ratio at week 4 (D) and week 10 (E). Quantification of HOPX/DAPI ratio at week 10 (K). Quantification of week 4 and 10 deep layer neuron proportion (F, G, H, I) and upper layer neuron proportion (L, M). One-way anova with Sidak’s multiple comparisons was used to calculate statistics for D-I, K. For L and M, unpaired t-test was used. *p<0.05, ** p<0.01, *** p<0.001, ns means not significant.

### *PTEN* p.I135L mutation and ASD genetic background dysregulate neurogenesis in cortical organoids

Since the cortical organoids produce heterogeneous cell types, we performed single cell RNA-sequencing (scRNA-seq) using the 10x platform on week 10 and 21 cortical organoids made from each of the isogenic ASD and control background lines. Three individual organoids from each genotype at each timepoint were mixed to account for organoid to organoid variation. After quality control, we profiled 95,227 cells for 12 samples, averaging 55k reads per cell. To assist with cell type annotation, we integrated our brain organoid datasets at each timepoint with a published, comprehensively annotated human fetal brain scRNA-seq dataset^38^. Our organoid scRNA-seq data integrated well with the fetal brain dataset. Diverse NPC and neuronal subtypes were produced at both 10 (Figure 4A, B) and 21 (Figure 4C, D) weeks, including IPC, oRG, ventricular radial glia (vRG), truncated radial glia (tRG) and cycling progenitors, as well as deep layer excitatory neurons, upper layer excitatory neurons and interneurons (Figure 4A, C, Figure S6). Several unknown clusters do not overlap with the brain dataset, which are likely non-brain cells due to mis-differentiation, which has also been found in other organoid datasets^19,39–41^. The integrated dataset enabled us to select specific cell types to perform differential expression analysis using GSEA between genotypes to study the effect of the *PTEN* p.I135L mutation and ASD genetic background.

**Figure 4.**
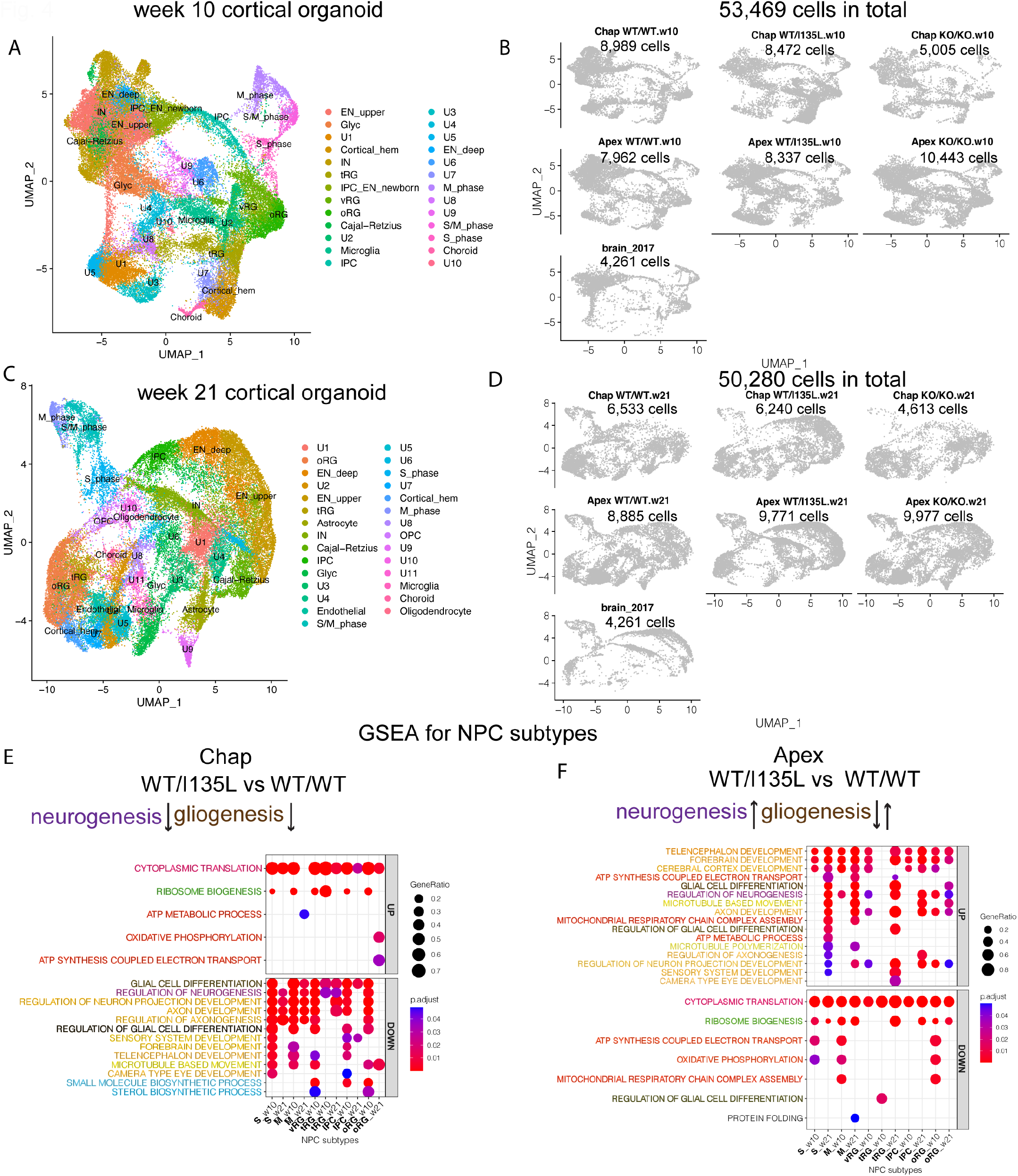
*PTEN* WT/I135L leads to dysregulated neurogenesis in NPC subtypes in cortical organoids. UMAP plots for visualizing clusters of NPC subtypes and neuron subtypes for week 10 (A) and week 21 cortical organoids (C). Split UMAP plots of cells for both week 10 (B) and week 21 organoids (D) demonstrate that scRNA-seq cell populations for each of the isogenic lines displayed similar spreads. (E) GSEA analysis on the effect of *PTEN* WT/I135L in the control genetic background in NPC subtypes from both week 10 and week 21 organoids. (F) GSEA analysis on the effect of *PTEN* WT/I135L in the ASD genetic background in NPC subtypes from both week 10 and week 21 dataset. P value was adjusted using Benjamini-Hochberg correction.

First, we performed GSEA analysis to study the effect of *PTEN* WT/I135L mutation in the control genetic background in NPC subtypes including IPC, oRG, vRG and tRG as well as cycling progenitor cells (S phase, M phase) at week 10 and week 21. The *PTEN* WT/I135L organoids displayed downregulated genes enriched for GO terms related with **neuron development** and **neurogenesis** at weeks 10 and 21, consistent with findings from NPC cultures (Figure 1B). Interestingly, the *PTEN* p.I135L mutation affected genes related to GO terms such as neuron development and neurogenesis in all NPC subtypes (Figure 4E, S6C). We identified additional downregulated genes enriched for GO terms such as “glia cell differentiation” and “regulation of gliogenesis”, suggesting that in addition to dysregulating neurogenesis, this *PTEN* mutation affected gliogenesis. Consistent with NPC cultures, we observed upregulated genes enriched for GO terms related with mitochondria including “ATP metabolic process” in M phase cycling progenitor and “oxidative phosphorylation” in oRG at week 21. Genes enriched for GO terms related with “ribosome biogenesis” were also upregulated as in 2D NPC cultures. Unexpectedly, we found that the *PTEN* p.I135L mutation downregulated genes enriched for GO terms related with biosynthetic processes in NPC subtypes including vRG, IPC and oRG in 3D organoid culture, whereas these GO terms enriched genes were upregulated in 2D NPC culture (Figure 1B and Figure 4E, S6C).

Second, we studied *PTEN* WT/I135L organoids on the ASD genetic background for the effects of the *PTEN* mutation on these NPC subtypes. The *PTEN* WT/I135L ASD organoids displayed upregulated genes enriched for GO terms related with **neuron development** and **neurogenesis** across NPC subtypes in both week 10 and week 21. We additionally identified dysregulated genes related with GO terms such as “glia cell differentiation”, “regulation of gliogenesis” and “regulation of glia cell differentiation”, which were upregulated in cycling progenitor cells, tRG and oRG at week 21, but were downregulated in tRG at week 10. These results suggested that *PTEN* p.I135L also dysregulates gliogenesis in the ASD genetic background. Consistent with 2D NPC findings, we observed microtubule related GO terms were enriched by genes upregulated in IPCs, oRGs as well as cycling progenitor cells. However, GO terms related with ribosome biogenesis and cytoplasmic translation were consistently enriched by genes that were downregulated by the *PTEN* mutation in the ASD genetic background, concordant with 2D NPC bulk RNA-seq findings (Figure 1C, Figure 4F, S6C).

Third, we asked whether ASD genetic background exerted similar effects on NPC subtypes in 3D culture as in 2D NPC culture. In the *PTEN* WT/WT genotype, the ASD genetic background resulted in downregulated genes enriched for GO terms related with **neurogenesis** and **gliogenesis** irrespective of NPC subtype, which is consistent with NPC 2D RNA-seq findings (Figure 2), Of note, genes that are associated with GO terms related to **neuron development** were both up and downregulated from the ASD genetic background at week 21 in oRG, whereas uniform downregulation effect was observed at week 10 for this cell type. ASD genetic background led to upregulated genes enriched for GO terms including “cytoplasmic translation”, “ribosome biogenesis”, “embryonic organ morphogenesis” and “mitochondria translation”, all terms were identified in 2D NPC culture (Figure 2A, Figure 5A, S6D). In the *PTEN* WT/I135L genotype, the ASD genetic background resulted in upregulated genes enriched for GO terms related to **neuron development, neurogenesis**, and **gliogenesis** in NPC subtypes, completely consistent with NPC bulk RNA-seq results (Figure 2B, Figure 5B, S6D). Interestingly, we observed that the ASD genetic background displayed downregulated genes enriched for GO terms related with ribosome biogenesis, cytoplasmic translation, mitochondria at week 10. However, these GO terms were activated at week 21 (Figure 5B, S6D). Contrary to the effect of ASD genetic background on the *PTEN* WT/I135L genotype, we observed uniform upregulated genes enriched for GO terms related to ribosome biogenesis, cytoplasmic translation, as well as uniform downregulation of genes associated with mitochondria related GO terms in NPC populations in organoids with the *PTEN* KO/KO genotype. In addition, the effect of ASD genetic background on neuron development and neurogenesis became asynchronized across NPC subtypes (Figure 2C and Figure 5C, S6D).

**Figure 5.**
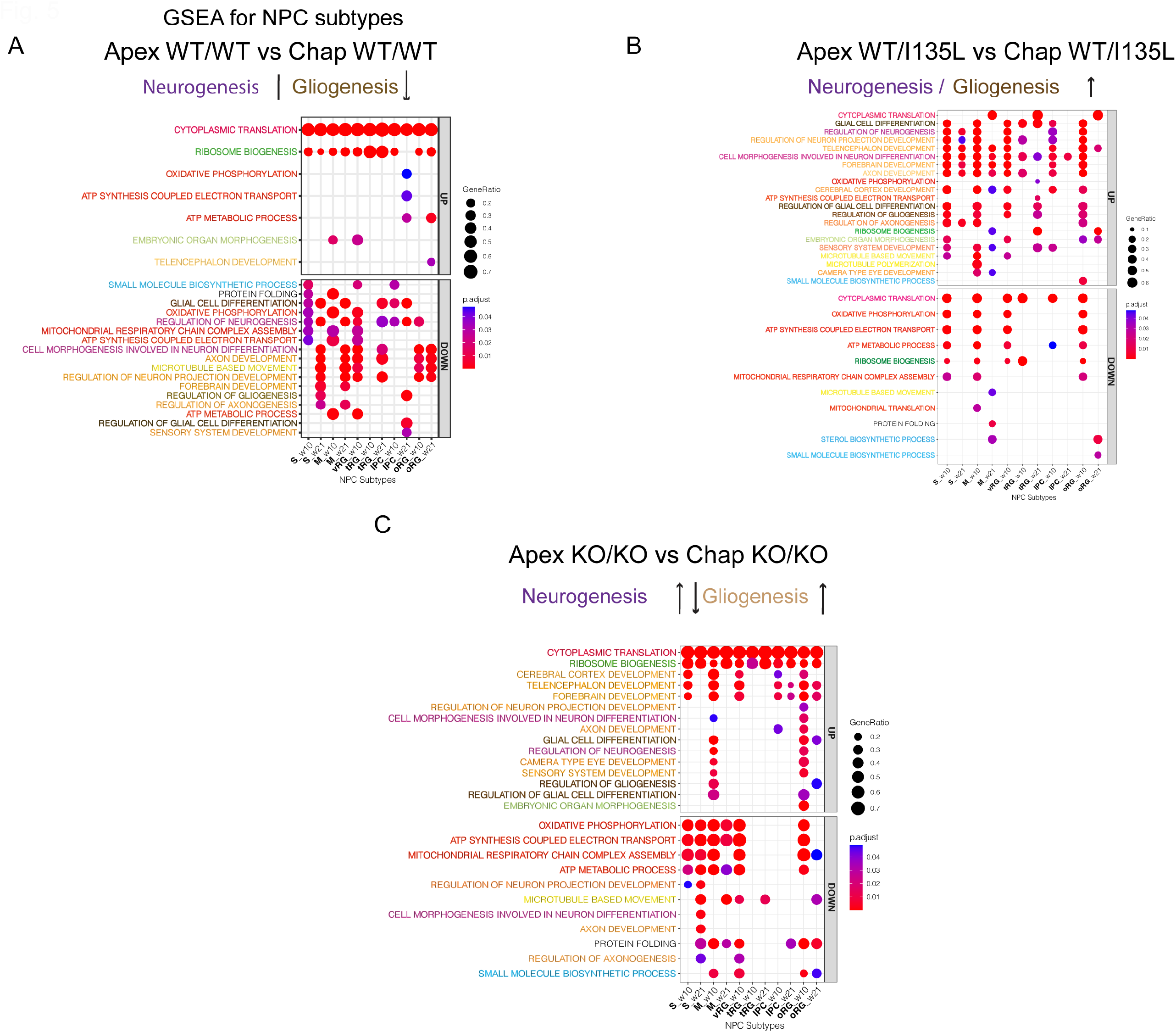
ASD genetic background dysregulates genes important for neurogenesis in NPC subtypes in cortical organoids. GSEA analysis for the effect of ASD genetic background in NPC subtypes with *PTEN* WT/WT genotype (A), with *PTEN* WT/I135L genotype (B) and with *PTEN* KO/KO genotype (C). GO terms were preselected for visualization and p values were adjusted using Benjamini-Hochberg correction.

These results provide additional evidence that the *PTEN* p.I135L mutation found in an ASD patient and ASD genetic background affect similar pathways, including dysregulating neurogenesis and gliogenesis. The ASD genetic background can reverse the effect of a mild ASD-specific *PTEN* WT/I135L mutation, while the effect of the ASD genetic background can also be modulated by the more severe complete loss-of-function of *PTEN*.

### *PTEN* p.I135L mutation accelerates the neuronal maturation of upper layer neurons in ASD genetic background

To determine whether the *PTEN* WT/I135L mutation affects neuronal maturation, we performed pseudotime analysis using Monocle3^42^ on the week 10 and week 21 scRNA-seq datasets. We used SOX2^+^ radial glia cells as the root cells to plot the pseudotime trajectory (Figure S7A-F), then plotted the distribution of deep layer excitatory neurons and upper layer excitatory neurons based on developmental pseudotime. At both week 10 and week 21, the *PTEN* WT/I135L organoids displayed accelerated upper layer neuron maturation only in the ASD genetic background (Figure 6B, D). Interestingly, *PTEN* WT/I135L organoids displayed decelerated upper layer neuron maturation in the control genetic background at week 21 but not at week 10. Deep layer neuron maturation remained unaffected by the *PTEN* p.I135L mutation in both control and ASD genetic background (Figure 6A, C). These results suggest that *PTEN* p.I135L accelerated upper layer neuron maturation in concert with increased NPC subtype proliferation, leading to overproduction of more mature upper layer neurons in the ASD genetic background.

**Figure 6.**
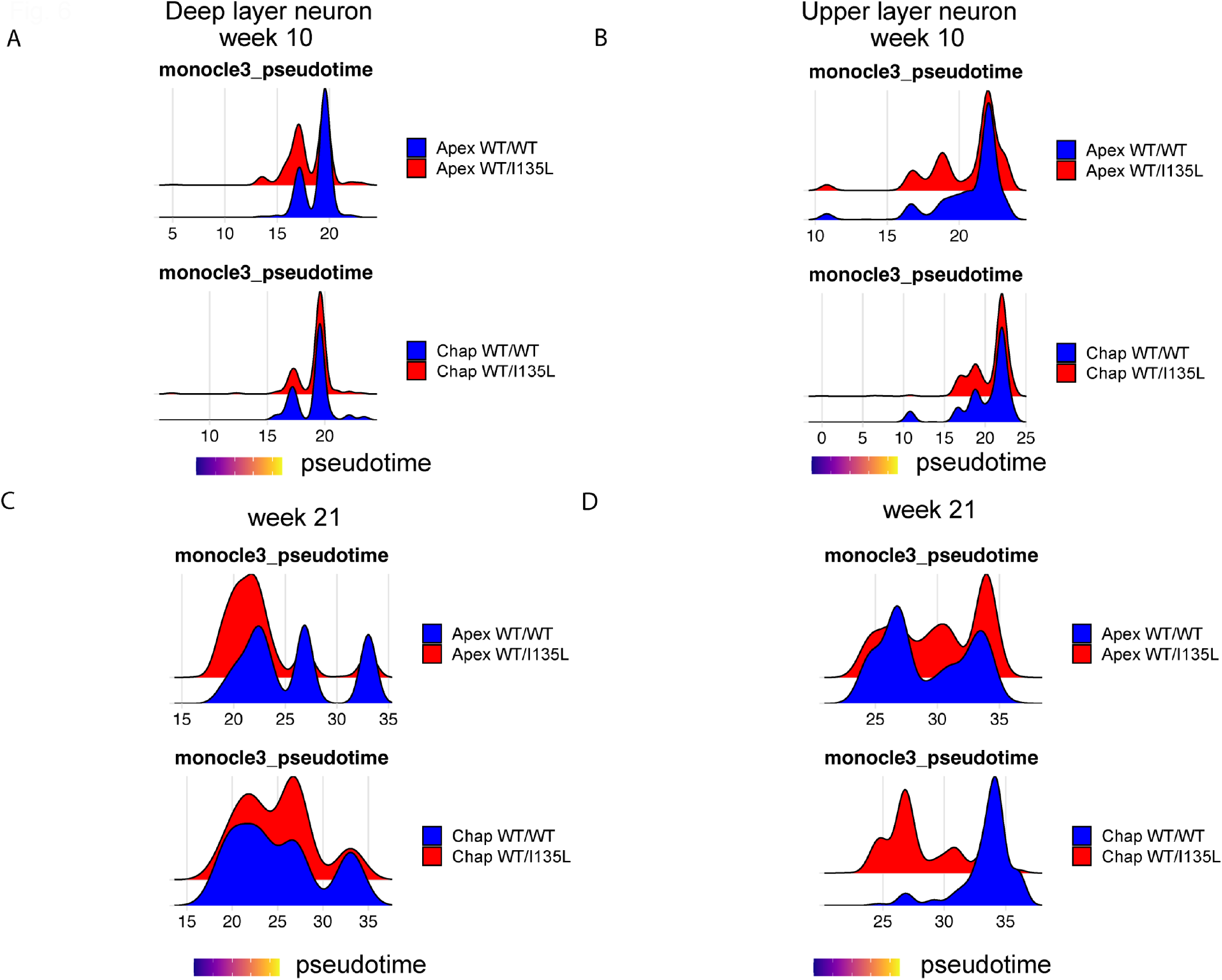
*PTEN* WT/I135L mutation accelerates the maturation of the upper layer neuron in the ASD genetic background. (A, B) Ridgeplot for visualization of the distribution of deep and upper layer neurons along the pseudotime at week 10. (C, D) Ridgeplot for visualization of the distribution of deep and upper layer neurons along the pseudotime at week 21.

### *PTEN* p.I135L mutation and ASD genetic background dysregulate synaptic transmission

To examine whether *PTEN* WT/I135L and ASD genetic background affected genes important for neuronal function in addition to accelerating neuronal maturation, we performed GSEA analysis on deep and upper layer excitatory neurons as well as interneuron subtypes. Neurons from the *PTEN* WT/I135L control background organoids displayed downregulated genes enriched for GO terms related with **synaptic function** (synaptic signaling, regulation of trans synaptic signaling and regulation of synaptic plasticity, Figure 7A). We also found that genes important for **action potential** were downregulated in *PTEN* WT/I135L control background organoids in both deep and upper layer excitatory neurons as well as interneurons. Genes enriched for GO terms important for RNA/protein expression such as “cytoplasmic translation,” “ribosome biogenesis” as well as “ATP metabolic process” were upregulated by the *PTEN* mutation in the control genetic background (Figure 7A).

**Figure 7.**
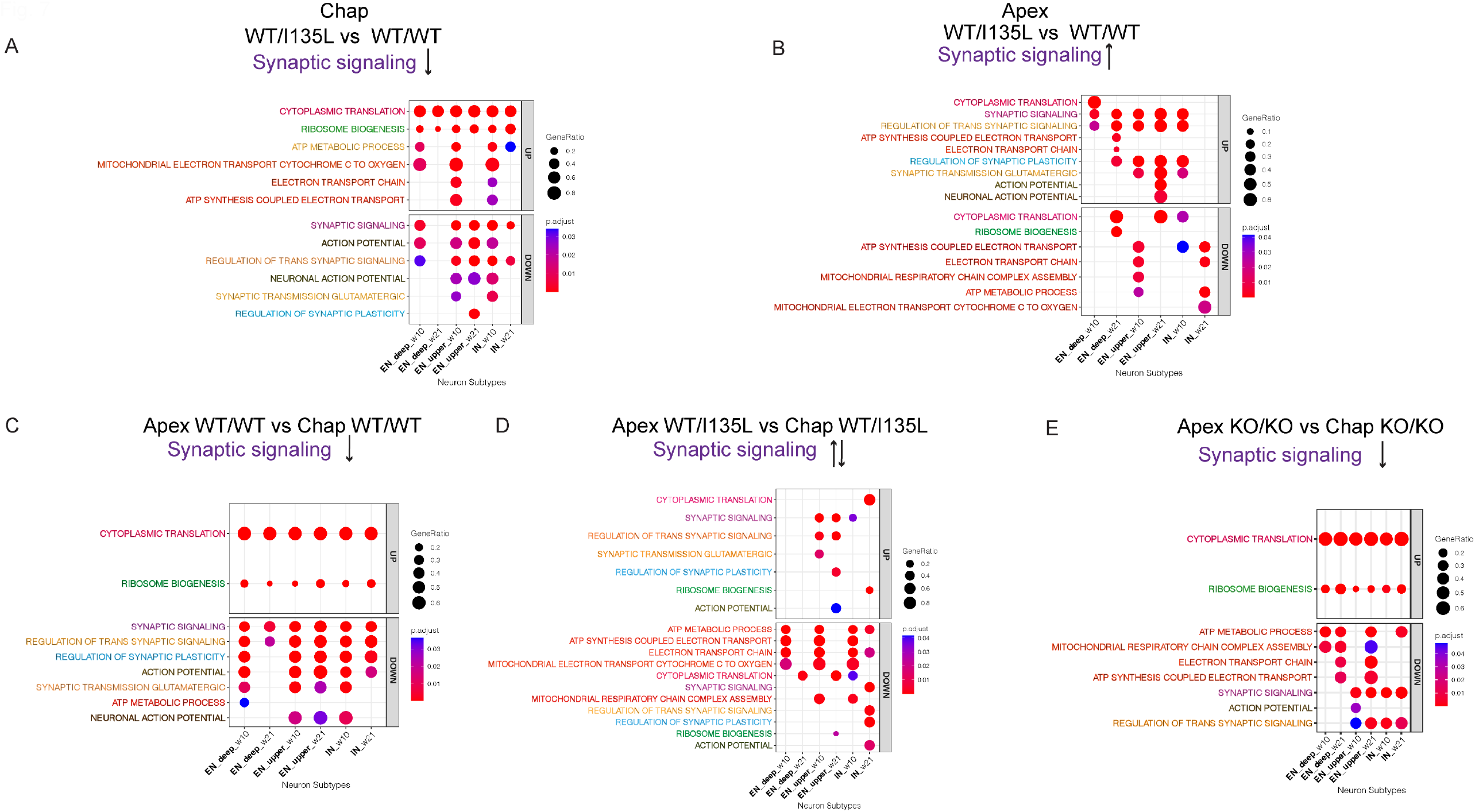
*PTEN* WT/I135L mutation and ASD genetic background dysregulate synaptic transmission. (A, B) GSEA analysis for the effect of *PTEN* WT/I135L mutation in control genetic background and ASD genetic background in different neuronal subtypes. (C, D, E) GSEA analysis for the effect of ASD genetic background in different neuronal subtypes with *PTEN* WT/WT genotype, with *PTEN* WT/I135L genotype, and with *PTEN* KO/KO genotype.

By contrast, in the ASD genetic background, genes enriched for GO terms related with **synaptic function** (synaptic signaling, regulation of trans synaptic signaling and regulation of synaptic plasticity) were all upregulated in *PTEN* WT/I135L organoids in deep layer and upper layer neurons as well as interneurons, which is the opposite direction of the effect of the same *PTEN* mutation on the control genetic background (Figure 7A, B, S6E). GO terms related with **action potential** were also enriched by genes upregulated by the PTEN WT/I135L mutation in the ASD genetic background, but only in upper layer neurons at week 21. Interestingly, we found that GO terms related with “cytoplasmic translation” were enriched by genes upregulated at week 10 in deep layer neurons, but were enriched by downregulated genes at week 21, whereas GO terms related with mitochondria (electron transport chain, ATP synthesis electron transport chain) was enriched by genes downregulated in upper layer excitatory neurons and interneurons, but was enriched by upregulated genes in the deep layer excitatory neurons (Figure 7B). All these results highlight the spatial and temporal regulation of gene expression in mature neuronal populations by *PTEN* WT/I135L in the ASD genetic background.

The reversed effect of the ASD *PTEN* mutation on **synaptic function** in the ASD genetic background versus its effect on the control genetic background motivated us to ask whether the ASD genetic background had a strong effect on neuronal GO terms. In the *PTEN* WT/WT genotype, neurons from organoids with the ASD genetic background displayed downregulated genes enriched for GO terms related with **synaptic function** (synaptic signaling, regulation of trans synaptic signaling, regulation of synaptic plasticity and action potential) irrespective of neuronal subtypes (Figure 7C), suggesting that the ASD genetic background indeed affects genes important for neuronal function. Neurons from organoids with the ASD genetic background displayed upregulated genes enriched for GO terms related with cytoplasmic translation and ribosome biogenesis. In the *PTEN* WT/I135L genotype, neurons from organoids in the ASD genetic background activated genes associated with GO terms related with **synaptic function** in the excitatory upper layer neurons for both week 10 and week 21 but not in deep layer excitatory neurons. In addition, we also observed a temporal effect of the ASD background on genes enriched for synaptic signaling in interneurons from organoids, being activated at week 10 and downregulated at week 21 (Figure 7D). In the *PTEN* KO/KO genotype, ASD genetic background downregulated genes enriched for GO terms related with **synaptic function** in both upper layer neurons and interneurons, but not deep layer neurons (Figure 7E).

Thus, similar to what we found in NPC subclasses in organoid models, genes enriched for neuronal GO terms such as **synaptic function** are dysregulated in a spatial and temporal dependent interplay between the ASD *PTEN* p.I135L mutation and the ASD genetic background.

## DISCUSSION

ASDs are heterogeneous genetic disorders that result from the complex interplay between genetic and environmental factors^43^. The number of identified ASD risk genes has grown over the past decade, especially those with high heritability and large effect size. One such gene, *PTEN*, is a major contributor to ASD risk, especially in the 20% of ASD individuals with macrocephaly^5,6^. Although *PTEN* is a well-known tumor suppressor, approximately 15% of ASD individuals with early brain overgrowth possess mutations in the *PTEN* gene^7^. The role of patient genetic background effect in modifying the risk of ASD has been more challenging to study, and experimental evidence for the role of genetic background in ASD development remains limited. A recent study^44^ identified an asynchronous effect of ASD risk gene mutations on neuron production by modeling the same ASD risk gene mutation on different control genetic backgrounds in cortical organoids, providing evidence that genetic background is an important factor when studying the role of a specific ASD risk gene disruption. Here, we have directly studied the cellular impact of a *PTEN* mutation found in an ASD individual in both control and ASD backgrounds by performing bi-directional CRISPR-Cas9 genome editing. An ASD patient-derived iPSC line possessing a point mutation of *PTEN* was used to produce an isogenic line with the correction of the *PTEN* mutation, both in the ASD background. Similarly, a control iPSC and the matched line with the induction of the same *PTEN* mutation were both in the control background. These isogenic *PTEN* iPSC panels enabled us to study the effect of the same ASD mutation in both control and ASD genetic background, but also the effect of ASD genetic background with different *PTEN* genotypes.

Our studies demonstrated that both the heterozygous *PTEN* p.I135L mutation and an ASD genetic background dysregulated genes important for cellular features of neurogenesis in induced, stable NPCs across three different passages, and in self-organizing cortical organoids across NPC subtypes at both week 10 and week 21. The *PTEN* p.I135L mutation resulted in increased proliferation in stable NPCs in both control and ASD backgrounds. However, the genetic backgrounds buffered the effect of the *PTEN* p.I135L mutation differentially. For example, comparing the effect of the same heterozygous *PTEN* p.I135L mutation in both control and ASD genetic backgrounds, we observed reversed effects of *PTEN* p.I135L on general GO terms from gene expression profiles related with neuron development and neurogenesis. The ASD genetic background also displayed profound effects dependent upon the *PTEN* genotype. In the *PTEN* WT/WT genotype, NPCs in the ASD background displayed more rapid proliferation compared to NPCs from the control genetic background, and the ASD genetic background resulted in the downregulation of gene expression profiles related to neurogenesis. However, in the *PTEN* WT/I135L genotype, there was a reversal of the effect of ASD genetic background on general GO terms from gene expression profiles related with neurogenesis, which was upregulated. In the *PTEN* KO/KO genotype, the effect of ASD genetic background on neurogenesis was non-uniform and asynchronous. Notably, most of these effects were entirely reproducible in three independent cell culture replicates over several passages in NPCs or in several organoids. These findings suggest that the *PTEN* mutations such as the heterozygous p.I135L found in ASD with minimal effects on canonical PTEN activity can be modified by the ASD genetic background to influence neurogenesis, while stronger *PTEN* mutations such as KO/KO are less sensitive to background modification of neurogenesis phenotypes, consistent with clinical findings in *PTEN*-related cancer-prone or ASD syndromes^10^.

Our study additionally identified, unexpectedly, that the *PTEN* p.I135L mutation impairs the canonical PTEN activity in the ASD genetic background but not in the control genetic background. Consistent with this biochemical readout, we also identified that the same *PTEN* mutation led to overproduction of NPC subtypes as well as neuronal subtypes only in the ASD genetic background, but not in the control genetic background in 3D cortical organoid model. Such an ASD genetic background dependent effect of the *PTEN* mutation on neurogenesis may be a direct effect of the genetic background on PTEN activity. We analyzed genetic variants (stop-gain and SNVs) and differentially expressed genes identified in the ASD line. Several ASD risk genes including *MET, FGF14, NTRK1, NTRK2, ITGB3, LAMB1, RELN* as well as *PRKCA* were identified which were all part of the PI3K/AKT pathway. These findings suggest one plausible molecular mechanism in which these additional dysregulated ASD risk genes in the genetic background may affect neurogenesis through modifying the effect of *PTEN* p.I135L mutation on the *PTEN* canonical activity.

The neuron overproduction finding in our *PTEN* panel cortical organoid model is also consistent with previous clinical findings, which identified 67% more neurons were present in the prefrontal cortex in postmortem brains from ASD individuals with brain overgrowth compared to healthy controls^45^. Our studies further indicated that ASD *PTEN* p.I135L mutation led to this neuron overproduction by both accelerating neuronal maturation and increasing NPC subtype proliferation only on the ASD genetic background. This clearly indicates the advantages of studying ASD candidate gene mutations in ASD genetic backgrounds using cortical organoid models.

In conclusion, our study provides direct evidence that ASD genetic background contributes to ASD pathology by working in concert with the ASD risk gene that displays a mutation found in ASD patients. Our study demonstrates that ASD genetic background modifies the effect of the *PTEN* p.I135L mutation, including on its *PTEN* canonical activity, resulting in accelerated neural maturation and elevated IPC and oRG production, which ultimately leads to the overproduction of the neuronal subtypes including deep and upper layer neurons in an ASD background-dependent manner in cortical organoids. By transcriptomic profiling the isogenic *PTEN* panel cortical organoids using scRNA-seq, our study revealed dysregulated gene expression profiles related with neurogenesis and gliogenesis in NPC subtypes, consistent with findings from bulk RNA-seq in 2D NPCs. We also uncovered dysregulated expression profiles related to synaptic function in neuronal subtypes resulted from both the *PTEN* p.I135L mutation and the ASD genetic background. Of note, our study revealed that both *PTEN* p.I135L mutation and an ASD genetic background reproducibly affected transcriptomic profiles related to both neurogenesis and gliogenesis in NPC subtypes. It is possible that the timing for the neurogenesis to gliogenesis switch is also impacted. A recent study^46^ identified that EGFR^+^ cells dramatically increased after gestational week 20, marking the onset of human gliogenesis, which was within the timing of the organoid models studied here. Future work on teasing out the effect of the ASD risk gene and ASD genetic background on the neuron to glia switch will yield additional insights on their roles in ASD pathology.

## ACKNOWLEDGMENTS

We would like to thank Drs. Ashleigh Schaffer, Helen Miranda and Charis Eng for their comments on the manuscript, and Dr. Ya Chen for technical assistance. This work was funded by an R01 from the NIH and NIMH (MH114601).

## AUTHOR CONTRIBUTIONS

S.F. and A.W.B. designed the research. S.F., L.A.B. and J.E. performed the research. S.F., L.A.B., J.E. and A.W.B. analyzed data. S.F. and A.W.B. wrote the paper.

## Supplemental information

## RESOURCE AVAILABILITY

### Lead contact

Please reach out to Dr. Anthony Wynshaw-Boris at if you want to request materials used in this study.

### Materials availability

Unique materials generated in this study will be fulfilled with a completed Materials Transfer Agreement.

### Data and code availability

Bulk RNA-seq and scRNA-seq raw data have been deposited at GEO and will be made publicly available as of the date of publication. All original code for scRNA analysis for this paper has been deposited and is publicly available under the following link: https://github.com/shuaifu93/codes-for-PTEN-manuscript

## EXPERIMENTAL MODEL AND SUBJECT DETAILS

### Human iPS cells

One control iPSC Chap and one ASD iPSC Apex described in our previous paper^16^ were used as the parental cell lines for generation of the isogenic *PTEN* iPSC panels. We introduced the ASD *PTEN* WT/135L mutation into clones of Chap iPSC using CRISPR-Cas9 genome editing^21^. This editing process also generated *PTEN* KO/KO clones in Chap. Additionally, we corrected the ASD *PTEN* WT/I135L mutation in Apex to WT/WT, and similarly generated Apex *PTEN* KO/KO. All cell lines have been karyotyped at NanoString using Human Karyotype Panel or at Cell Line Genetics using G-banding. No chromosome abnormality was observed in our cell lines. All iPSCs were maintained in mTeSR plus medium (StemCell Technologies) using a feeder-free culture protocol in six-well plates coated with Vitronectin (Gibco, A31804) ^47^. iPSCs were cultured at 37°C and 5% CO_2_ with feeding of 2 mL mTeSR plus per well every other day. Passaging of iPSC colonies was carried out using 0.5mM EDTA (Thermo Fisher).

## METHOD DETAILS

### Generation of isogenic iPSCs using CRISPR-mediated genome editing

We amplified a 1.6kb PCR product from Apex genomic DNA spanning the heterozygous *PTEN* I135L mutation site. This product, containing both Wildtype and Mutant *PTEN* fragments with and without the point mutation, was fused with the HindIII digested pUC19 vector via Gibson Assembly^48^ and transformed into competent cells to produce the donor plasmids. Donor plasmid identity was confirmed via Sanger sequencing for the plasmids derived from single colonies to determine whether the clone contained the Wildtype *PTEN* donor plasmid, which was used for CRISPR correcting mutation in ASD iPSC, or the Mutant *PTEN* donor plasmid, which was used for CRISPR installing *PTEN* I135L mutation. We co-transfected plasmids (100ng) containing double nickase cas9 (Cas9n) and two guide RNAs (guide RNA #1 TGTCATCTTCACTTAGCCAT, guide RNA #2 AAAGCTGGAAAGGGACGAAC) to target Cas9n to the *PTEN* locus, along with the appropriate donor plasmid (400ng) using Lipofectamine™ Stem Transfection Reagent (Invitrogen, STEM00001) in E8 medium in one well of a 24-well plate. After 48h of transfection, iPSCs were dissociated into single cells using TrypLE™ Express Enzyme (Gibco, 12605093) which were then seeded into 6 well plates at 250 cells per well. We then picked single colonies into individual wells of 48 well plates for further expansion and Sanger sequencing. Sanger positive clones were then subjected to targeted amplicon sequencing using 2×250bp Miseq to ensure purity of clones and that no unintentional indels introduced in the edited clones had occurred. Fastq files from Miseq were processed with CRISPResso (version: 2.0.29) for CRISPR-edit analysis^49^.

### Generation of dorsal cortical NPC

We differentiated the isogenic iPSCs from the control and ASD *PTEN* panels into dorsal NPCs using the PSC Neural Induction Medium protocol (A1647801, Gibco). Briefly, iPSCs were seeded in 6 well plates at approximately 20% confluence. On the second day, the media was replaced with neuronal induction medium (NIM), which was used for 7 days with 3 ml media changes every other day. At day 7, cells were dissociated into single cells using Accutase (Stemcell Technologies, 07922) and were seeded at 10^6^ cells per well in neural expansion medium (NEM) in 6 well plates coated with Geltrex (A1413201, Gibco). This was considered as Passage 0 (P0). NPCs were passaged when reaching 85% confluence.

### Immunofluorescence staining for 2D NPC

NPCs were washed in PBS, fixed with 4% paraformaldehyde in PBS for 15 min at room temperature, washed 3 times with PBS, each for 5 min, and permeabilized with 0.5% Triton X-100 in PBS for 10 min. Blocking was performed with 0.1% Triton X and 1% BSA in PBS for at least 1 h, followed by overnight primary antibody incubation at 4°C. The following primary antibodies were diluted in the blocking solution: FOXG1 (Abcam, 1:500) and PAX6 (BD Biosciences; 1:500). Following three 5 min washes in PBS, the cells were incubated with the appropriate secondary antibodies conjugated with Alexa Fluor 488 or Alexa Fluor 555 (Thermo Fisher Scientific; 1:500) for 30 min at room temperature, followed by three 5 min washes in PBS. Cells were counterstained with DAPI in PBS (Sigma Aldrich, 1:2000) for 5 min, then washed 3 times. Fluorescence was visualized with the Leica DM6000 inverted microscope in the CWRU Light Microscopy Imaging Core (NIH Grant S10-RR021228). Images were acquired using the Q-Imaging Retiga Xi Firewire High-Speed, 12-bit cooled CCD camera and Volocity software.

### Western blotting

When NPCs reached 85% confluences in 6 well plates, medium was aspirated, cold PBS was added, cells were scraped off the plate, and resuspended in Mammalian Protein Expression Reagent (M-PER, 78501, Thermo Scientific) supplemented with Halt Protease and Phosphatase Inhibitor Single-Use Cocktail (78442, Thermo Scientific). Protein lysate quantification was done using the Bradford assay (Bio-Rad). 10 μg protein lysates were loaded into a NuPAGE 4%-12% 17-well gel (NP0329BOX, Invitrogen), and transferred onto nitrocellulose membranes. Blots were blocked in Intercept (TBS) Blocking Buffer for 1h, primary antibodies (PTEN, 9188S; p-AKT S473, 4060S; p-AKT T308, 2965S; Total AKT, 2920S from Cell Signaling Technology, Beta Actin sc-47778, Santa Cruz Biotechnology) were incubated in TBS blocking buffer overnight (1:1000 dilution), then blots were washed and incubated with fluorescent secondary Antibody (1:20,000) for 1h. Blot images were captured and quantified using Odyssey XF Imaging System.

### Population doubling time

NPCs at passage 3 were seeded at a density of 7×10^4^/cm^2^ in four wells of the 6 well plate pre-coated with Geltrex. When NPC wells reached 85% confluence, cells were dissociated with Accutase and cells from three wells were counted. Doubling time was calculated based on culture time and the ratio of cell number before seeding and after harvesting. Cells in the fourth well of the 6-well plate were collected for Bulk RNA-seq. Similar procedures were done continuously until P10. Proliferation assays were done simultaneously for all isogenic clones in the control and ASD backgrounds. Three separate proliferation assays were done for the isogenic clone sets.

### Bulk RNA sequencing

#### RNA extraction

NPCs were collected by adding 1ml Trizol into one well of a 6 well plate washed with DPBS, and stored at -80°C from P3, P4, P5 across 3 batches. RNA was then extracted in a single batch using the Direct-zol RNA Miniprep Plus Kit (Zymo Research). The extracted RNA was subjected to 150bp unstranded paired-end poly-A RNA sequencing on the Illumina platform Novaseq (Novogene).

#### Data alignment and analysis

Paired-end RNA sequencing reads were aligned to the human reference transcriptome (Ensembl version 103) using Kallisto version 0.46.2^50^ for transcript abundance quantification. All downstream analysis was performed using RStudio (2022.07.1, build 554) with R version 4.2.1. Tximport R package was used to summarize transcript quantification data to genes^51^ followed by the R package edgeR TMM method for normalization^52^. Genes with <1 CPM among 3 sample replicates, were filtered out. Variance-stabilization for the normalized filtered data was carried out with limma voom^53^, and differentially expressed genes were identified with linear modeling using limma with Benjamini-Hochberg multiple testing correction.

To perform Gene Set Enrichment Analysis (GSEA) analysis, we used the Molecular Signatures Database (MSigDB v7.5.1) and C5 subcluster “biological process” was used for Gene Ontology (GO) analysis. All gene lists that were used for the input were ranked based on the t-statistic of the limma voom output, and we performed the analysis in the R package ClusterProfiler 4.5.1 using default parameters (minGSsSize = 10, maxGSSize = 500). The “comparecluster” function was used to visualize the GO term enrichment from multiple passages at the same time^54^.

#### Genetic analysis for the ASD line

Whole exome sequencing for the ASD line Apex fibroblast was performed in our previous paper^16^. Stop gains and stop loss mutations as well as SNVs were pulled out from the original publication^16^, in total 68 genes with either stop gains or stop loss were found, and 380 genes were found to have SNVs in Apex.

#### Differential expression analysis for Apex vs Chap NPC

9 samples from Apex and 9 samples Chap NPC were used for differential expression analysis, decideTests function in the limma R package was used to identify the differentially expressed genes with cutoff adjusted p value = 0.05 and log2 fold change = 0.6 with Benjamini-Hochberg multiple testing correction. In total, we identified 1632 downregulated genes and 1729 upregulated genes Apex vs Chap.

### Cortical organoid production

We used published organoid culture protocols with modifications^14,35,36^. When iPSC cultures reached ∼80% confluency, the medium was aspirated and wells rinsed twice with PBS. 1 mL of Accutase was added per well of the 6-well plate and incubated for 2 minutes at 37°C, 5% CO_2_, then 2ml mTESR plus medium was added to the 6 well plate. Cells were then scraped off the plate. Gentle trituration was performed using a 5ml pipette to achieve a single cell suspension, which was transferred to a 15 mL conical tube, collected by centrifugation at 1200 rpm for 4min. Cells were resuspended in Cortex Differentiation Medium (Glasgow-MEM, 20% KSR, 1X NEAA, 1X Sodium Pyruvate, 1x Glutamax, 0.7% Beta-Mercaptoethanol (BME), 1% Pen/Strep) plus ALI (1μM **A**83-01 (Stem cell Technologies), 100nM **L**DN-193189(Sigma-Aldrich), 3μM **I**WR-1(Sigma-Aldrich)) plus 10 μM ROCK inhibitor Y-27632 (Tocris). Cells were counted using a hemocytometer, and seeded at 9,000 cells per well of a 96-well V-bottom ultra-low attachment plate (S-Bio, MS-9096VZ) in 100μl volume for 48 hours. On day 2, 100μl Cortex Differentiation Medium with ALI and ROCK inhibitor were added. From day 4 to day 10, 100μl of the old medium was aspirated and 100μl of fresh Cortex Differentiation Medium with ALI was added every other day. From day 10 to day 18, only AL was added to the Cortex Differentiation Medium. At day 18, organoids in 96 well plates were transferred to ultra-low attachment 6 well plates (Costar, #3471) in Early Organoid Medium (DMEM/F12, 1x N2, 1x Glutamax, 1x Chemically Defined Lipid Concentrate, 1% Pen/Strep and 0.1% Fungizone), with approximately 16 organoids per well and 3 wells for each genotype and cultured on an orbital shaker (Thermo Scientific) at 85 rpm at 37°C, 5% CO_2_. Medium changes were performed every other day. At day 35, the culture medium was switched to Late Organoid Medium (DMEM/F12, 10% FBS, 1x N2, 1x Glutamax, 1% Pen/Strep, 0.25 % Fungizone, 0.1% Heparin, 1x Chemically Defined Lipid Concentrate) supplemented with fresh 1% Matrigel. From day 70 onwards, culture medium was switched to Final Organoid Medium (DMEM/F12, 10% FBS, 1x N2, 1x B27 without vitamin A, 1% Pen/Strep, 0.25 % Fungizone, 0.1% Heparin, 1x Chemically Defined Lipid Concentrate) with 2% Matrigel added fresh, and medium was changed every other day.

### Organoid size analysis

Bright field images of organoids at week 4 and week 8 were obtained via the Leica DMi1 Inverted Microscopes, and the organoid size determined using a pixel-based method with customized script using ImageJ version 1.53k.

### Fixation and frozen sectioning of organoids

Organoids were fixed on ice in 4% paraformaldehyde for 30min, followed by three rinses in PBS, and allowed to sink in 30% sucrose in PBS at 4° C overnight. Organoids were then placed in cryomolds (Tissue Tek, 4565) with OCT compound (Tissue Tek, 4583) and 30% sucrose (1:1), snap frozen on dry ice and stored at -80°C. Organoids were sectioned via cryostat (Leica) sequentially at 20 μm thickness. Sections were placed on microscope glass slides, dried overnight and stored at -20°C for subsequent immunohistochemistry.

### Immunohistochemistry of organoid frozen sections

Sections were thawed and air-dried at room temperature then washed two times in TBS-T (Tris Buffered Saline, 0.1% Triton X-100) for 10 minutes per wash. Slides were blocked with 10% donkey serum in TBS-T for 30-45 minutes at room temperature in a humidifying chamber. Slides were then incubated with primary antibodies diluted in blocking solution at 4° C overnight. The next day, slides underwent 4 washes in TBS-T (10 minutes per wash), then were incubated with secondary antibodies at room temperature for 1hr. Slides underwent another 3 washes in TBS-T (10 minutes per wash). Slides were counterstained with DAPI in TBS for 5min then washed in TBS for 5min before coverslips were mounted with fluoromount (Sigma, F4680). Tissue sections were imaged at a 20x objective on a Hamamatsu Nanozoomer S60 slide scanner in the CWRU Light Microscopy Imaging Core. Image analysis was performed on NPD.view2 (Windows (Ver. 2.9.29)) and ImageJ (version 1.53k).

### Dissociation of brain organoids and scRNA-seq

Organoids were dissociated into single-cell suspension with the Papain dissociation system (Worthington Biochemical, LK003150). Cells were then loaded onto the Chromium Next GEM Chip G Single Cell Kit, (10x Genomics, 1000120), and were processed with Chromium Controller to generate single-cell gel beads in emulsion. The Chromium Next GEM Single Cell 3’ Kit v3.1 (10x Genomics, 1000268) was used for all scRNA-seq library preparations. Libraries were pooled from different samples based on molar concentrations and sequenced on NovaSeq (Illumina) with 28 bases for read 1, 91 bases for read 2 (Novogene).

### scRNA-seq data analysis

#### Read alignment and processing

Raw demultiplexed fastq files were uploaded into the 10x genomics cloud, and reads were aligned to Human (GRCh38) 2020-A using Cell Ranger v6.1.2 with default settings. The generated “filtered cell matrix HDF5 file” for each sample was loaded into Seurat 4.1.1^55^. We removed all cells expressing less than 200 genes as well as cells containing more than 30% mitochondria content. SCTransform v2 was used to process each Seurat object to account for sequencing depth and batch effect^56^, then the top 3000 variable features were selected using the SelectIntegrationFeatures function. We integrated the organoid datasets collected from the same timepoint as well as fetal brain dataset^38^ using canonical correlation analysis (‘CCA’). Principal component analysis (PCA) was performed on the scaled data for the variable genes. PCA was then used to cluster cells using Seurat FindNeighbors function with reduced dimension 1:30, followed by FindClusters with resolution = 1.5. UMAP was used for dimension reduction and visualization. The top expressed genes in each cluster were derived by first normalizing SCT reads with PrepSCTFindMarkers and then using FindMarkers function in the “Seurat” R package. Cell types for each cluster were manually annotated based on the fetal brain dataset^38^, marker gene expression (Figure S6) and literature searches.

#### Differential expression analysis for scRNA-seq

We used the Seurat’s FindMarkers to perform differential expression between genotypes of interest using their default, the Wilcoxon test. In addition, we removed genes which are expressed in less than 10% of the cells using min.pct = 0.1. We filtered all the genes using logfc.threshold = -Inf, then ranked the genes for GSEA analysis based on avg_log2FC. GSEA between genotypes was performed using clusterprofiler 4.5.1, and the comparecluster function was used to visualize the GO term enrichment across different NPC subtypes or neuronal subtypes with default settings^54^.

#### Pseudotime analysis

We performed Monocle3 (version 1.2.9) pseudotime analysis^42^ on filtered datasets including cycling progenitor cells, vRG, tRG, oRG, IPC, Cajal-Retzius cells, deep layer excitatory neurons and upper layer excitatory neurons as well as newborn excitatory neurons. SOX2^+^ radial glia cells were set as the root cells. We down-sampled the cell number to ensure equal numbers between genotype comparisons, and we plotted the distribution of cells from deep and upper layer neuron clusters along the pseudotime trajectory using ridgeplots.

## QUANTIFICATION AND STATISTICAL ANALYSIS

### Organoid size analysis

In summary, for week 4, n = 45 for Chap *PTEN* WT/WT organoids, *n* = 42 for Chap *PTEN* WT/I135L organoids, *n* = 45 for Chap *PTEN* KO/KO organoids, n = 46 for Apex *PTEN* WT/WT organoids, n = 44 for Apex *PTEN* WT/I135L organoids and n = 21 for Apex *PTEN* KO/KO organoids. For week 8, n = 11 for Chap *PTEN* WT/WT organoids, n = 11 for Chap *PTEN* WT/I135L organoids, n = 12 for Chap *PTEN* KO/KO organoids, n = 10 for Apex *PTEN* WT/WT organoids, n = 9 for Apex *PTEN* WT/I135L organoids and n = 8 for Apex *PTEN* KO/KO organoids. *P* values were calculated using one-way ANOVA among the same genetic background and then adjusted using Sidak’s multiple correction.

### scRNA-seq analysis

In each dataset, three individual organoids were mixed for each genotype profiled. We applied cutoff nFeature_RNA > 200, and percent.mt < 30, and in total 12 datasets (each dataset contains a mixture of three organoids) that passed quality control were used in downstream analysis, with a total of 95,227 cells, averaging 55k reads per cell. P values for differential expression were calculated using the Wilcoxon rank sum test and p values for GSEA analysis were calculated using the GSEA function from clusterprofiler 4.5.1 R package with Benjamini-Hochberg correction.

### Bulk RNA-seq analysis

In each genotype, RNA from three independent cell culture experiments across three different passages including passage 3, passage 4, passage 5 were profiled, for a total of 54 samples. Libraries were sequenced with paired-end 150bp using Novaseq, with an average of 30M paired end reads per sample. P values for GSEA analysis were calculated using the GSEA function from clusterprofiler 4.5.1 R package with Benjamini-Hochberg correction.

### Immunohistochemistry

For week 10 HOPX/DAPI, SATB2/DAPI, TBR2/DAPI, CTIP2/DAPI, TBR1/DAPI ratio quantification, n = 5 for Chap *PTEN* WT/WT organoids, *n* = 3 for Chap *PTEN* WT/I135L organoids, *n* = 3 for Chap *PTEN* KO/KO organoids, n = 3 for Apex *PTEN* WT/WT organoids, n = 4 for Apex *PTEN* WT/I135L organoids and n = 1 for Apex *PTEN* KO/KO organoids. For week 4 TBR2/DAPI, CTIP2/DAPI, TBR1/DAPI radio quantification, n = 4 for Chap *PTEN* WT/WT organoids, n = 5 for Chap *PTEN* WT/I135L organoids, n = 5 for Chap *PTEN* KO/KO organoids, n = 5 for Apex *PTEN* WT/WT organoids, n = 4 for Apex *PTEN* WT/I135L organoids and n = 3 for Apex *PTEN* KO/KO organoids. In addition, for each organoid quantified, the average proportion value was used from three separate section quantification (distance between section 200 μm) for downstream statistical analysis. P values were calculated using one-way ANOVA and were then adjusted using Sidak’s multiple comparisons.

### Western blotting

Protein lysates from NPCs at P5, P6 and P7 for each genotype were used for the western blot. Two batches of P5 lysates were used.

**Figure S1.**
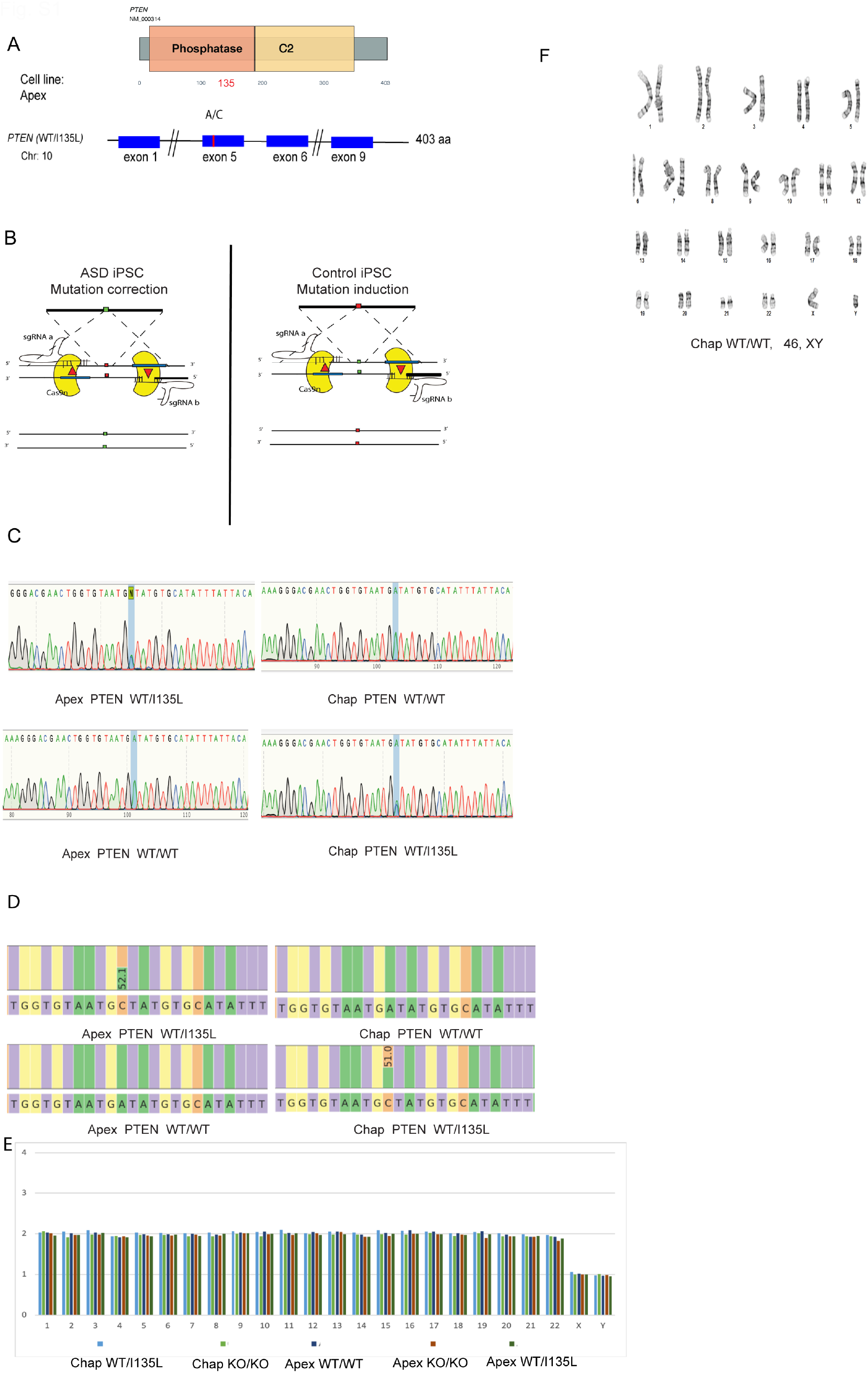
Generation and validation of isogenic PTEN panel iPSCs, related to Figure 1. (A) Diagram of WT/I135L mutation in *PTEN* locus. Mutation is located within the phosphatase domain. (B) Diagram of the strategy used for performing genome editing using dual Cas9n to generate isogenic induction line for control iPSC Chap, and to generate isogenic correction line from ASD iPSC Apex. (C) Sanger sequencing was used to verify the generated isogenic lines. (D) Targeted amplicon sequencing using Miseq (2x 250bp paired-end reads) for validation of CRISPR-edited clones. (E) Karyotyping for isogenic iPSC using the Nanostring human karyotype panel. (F) Karyotyping for control iPSC Chap using G-banding.

**Figure S2.**
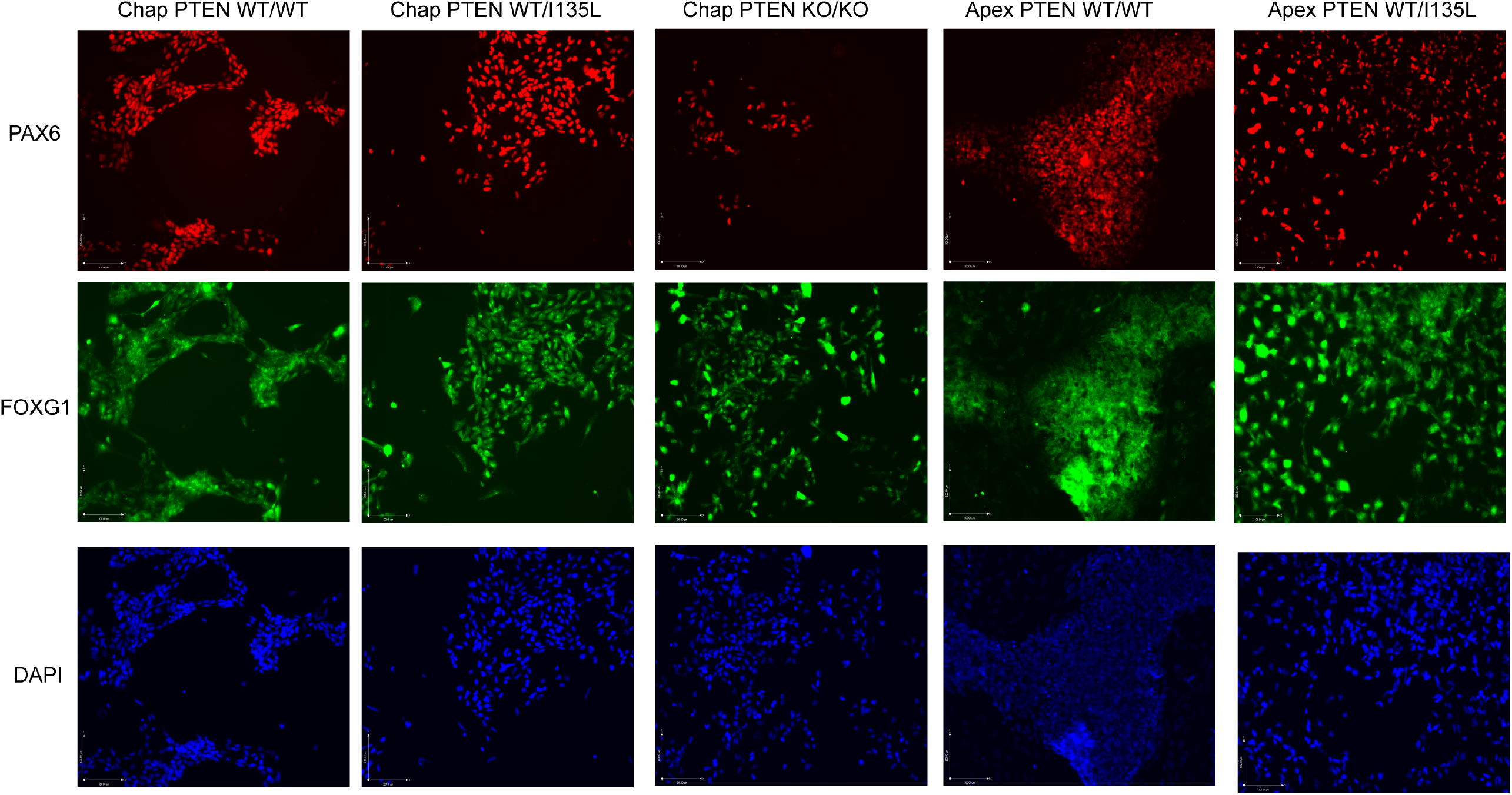
Immunofluorescence of 2D NPCs for isogenic *PTEN* panels, related to Figure 1. Representative images for staining of stable, 2D NPCs derived from isogenic *PTEN* iPSCs. NPCs at passage 6 were fixed and stained with anti-FOXG1 (1:500, ABCAM, ab18259) and anti-PAX6 (1:500, BD Biosciences, 561462) antibodies and counterstained with DAPI, scale bar = 100μm.

**Figure S3.**
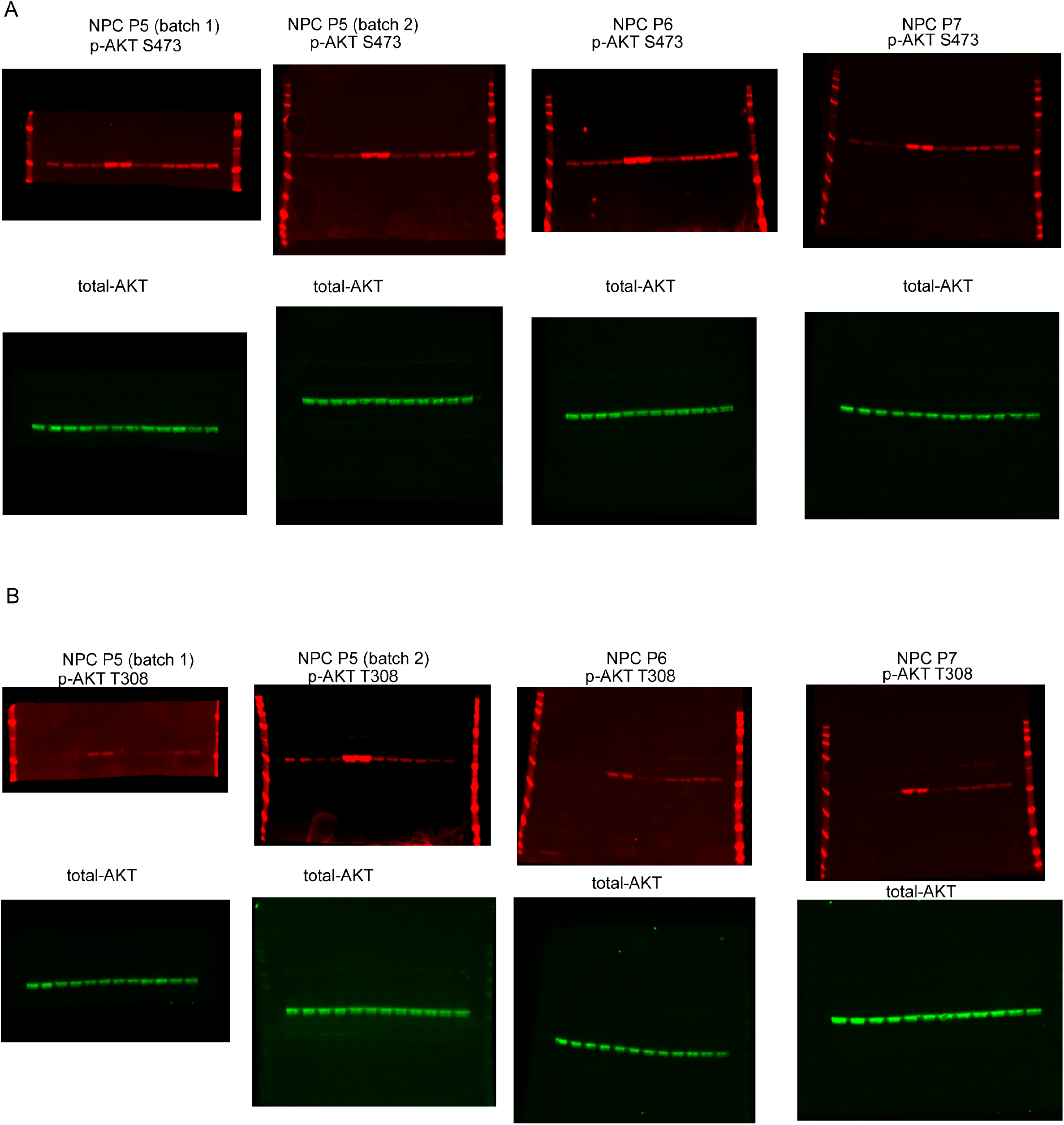
*PTEN* p.I135L mutation activates PI3K/AKT in ASD genetic background but not control genetic background, related to Figure 1. (A) Raw uncropped western blot images which were used for Figure 1E quantification of p-AKT S473 to total AKT ratio. (B) Raw uncropped western blot images which were used for Figure 1F quantification of p-AKT T308 to total AKT ratio.

**Figure S4.**
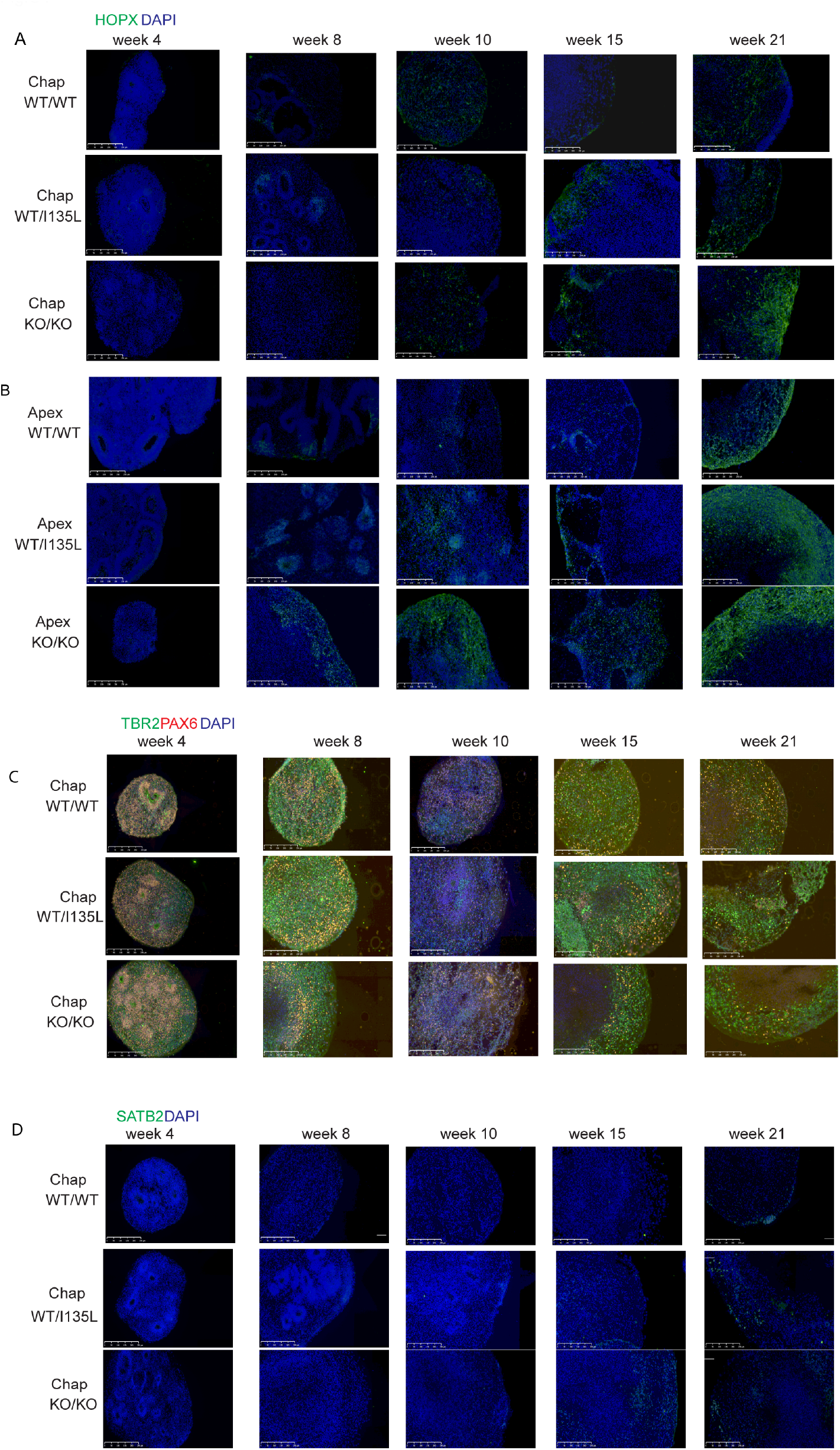
*PTEN* p.I135L mutation leads to overproduction of oRGs and IPCs in the ASD genetic background, related to Figure 3. Representative IHC images for HOPX and DAPI staining for control background (A) and ASD background organoids (B), scale bar = 250μm. (C) Representative IHC images for TBR2, PAX6 and DAPI staining. (D) Representative IHC images for SATB2 and DAPI staining for control background organoids, scale bar = 250μm.

**Figure S5.**
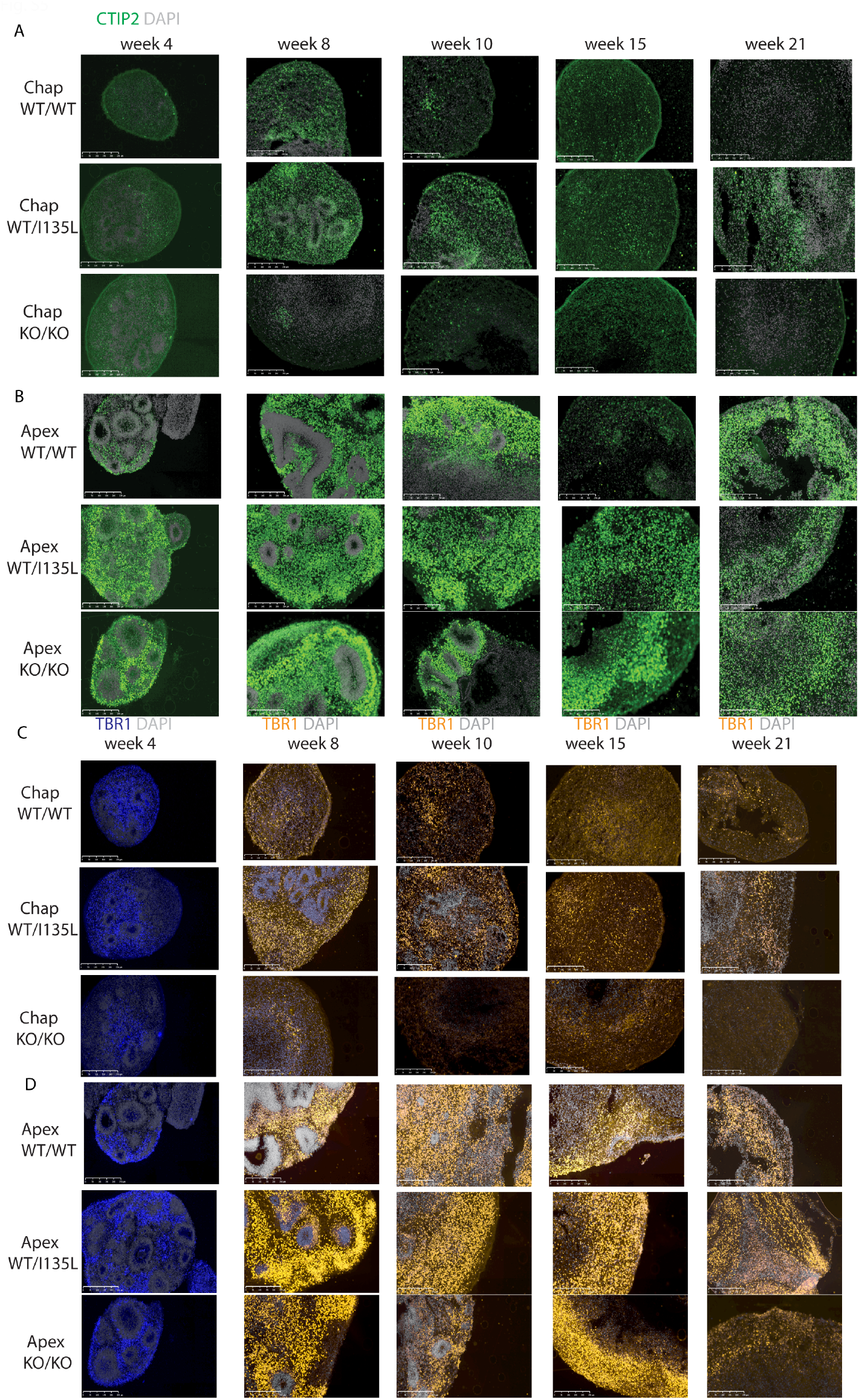
PTEN p.I135L mutation leads to overproduction of deep layer neurons in ASD genetic background, related to Figure 3. (A) Representative IHC images for CTIP2 and DAPI staining for control background organoids. (B) Representative IHC images for CTIP2 and DAPI for ASD background organoids. (C) Representative IHC images for TBR1 and DAPI staining for control background organoids. (D) Representative IHC images for TBR1 and DAPI staining for ASD background organoids. scale bar = 250μm.

**Figure S6.**
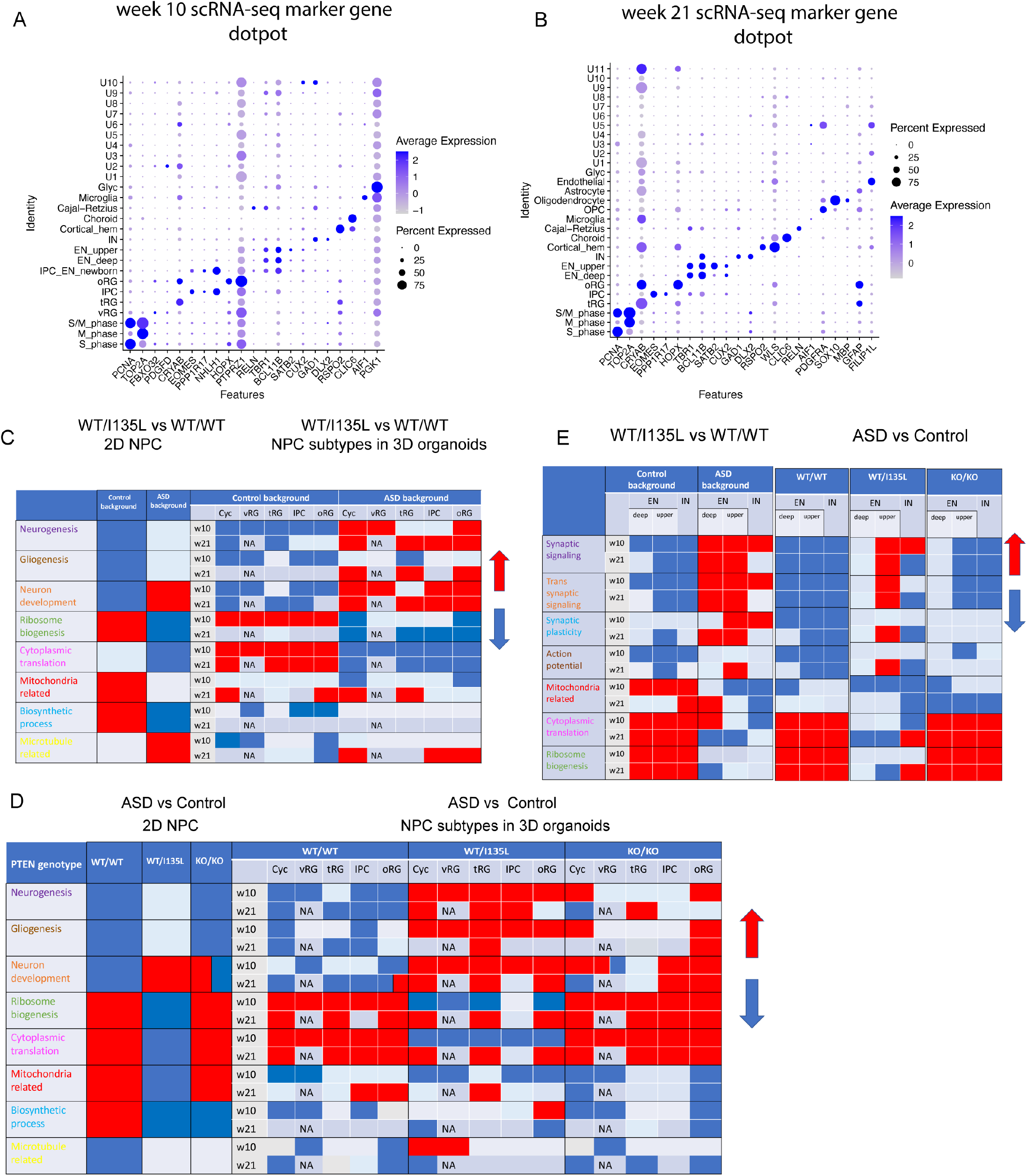
scRNA-seq marker genes expression and summary tables for GSEA results, related to Figure 4, 5 and Figure 7. Dotplot for visualizing marker gene expression for annotated cell types for both week 10 (A) and week 21 (B) integrated datasets. (C) Summary table for the GSEA results for studying the effect of PTEN WT/I135L in both control background and ASD background from both 2D NPC and NPC subtypes from 3D cortical organoids. (D) Summary table for the GSEA results for studying the effect of ASD genetic background in 2D NPC and NPC subtypes from 3D cortical organoids. (E) Summary table for the GSEA analysis for the effects of PTEN WT/I135L mutation in both control and ASD genetic background as well as the effects of ASD genetic background with *PTEN* WT/WT genotype, with *PTEN* WT/I135L genotype, and with *PTEN* KO/KO genotype in different neuronal subtypes.

**Figure S7.**
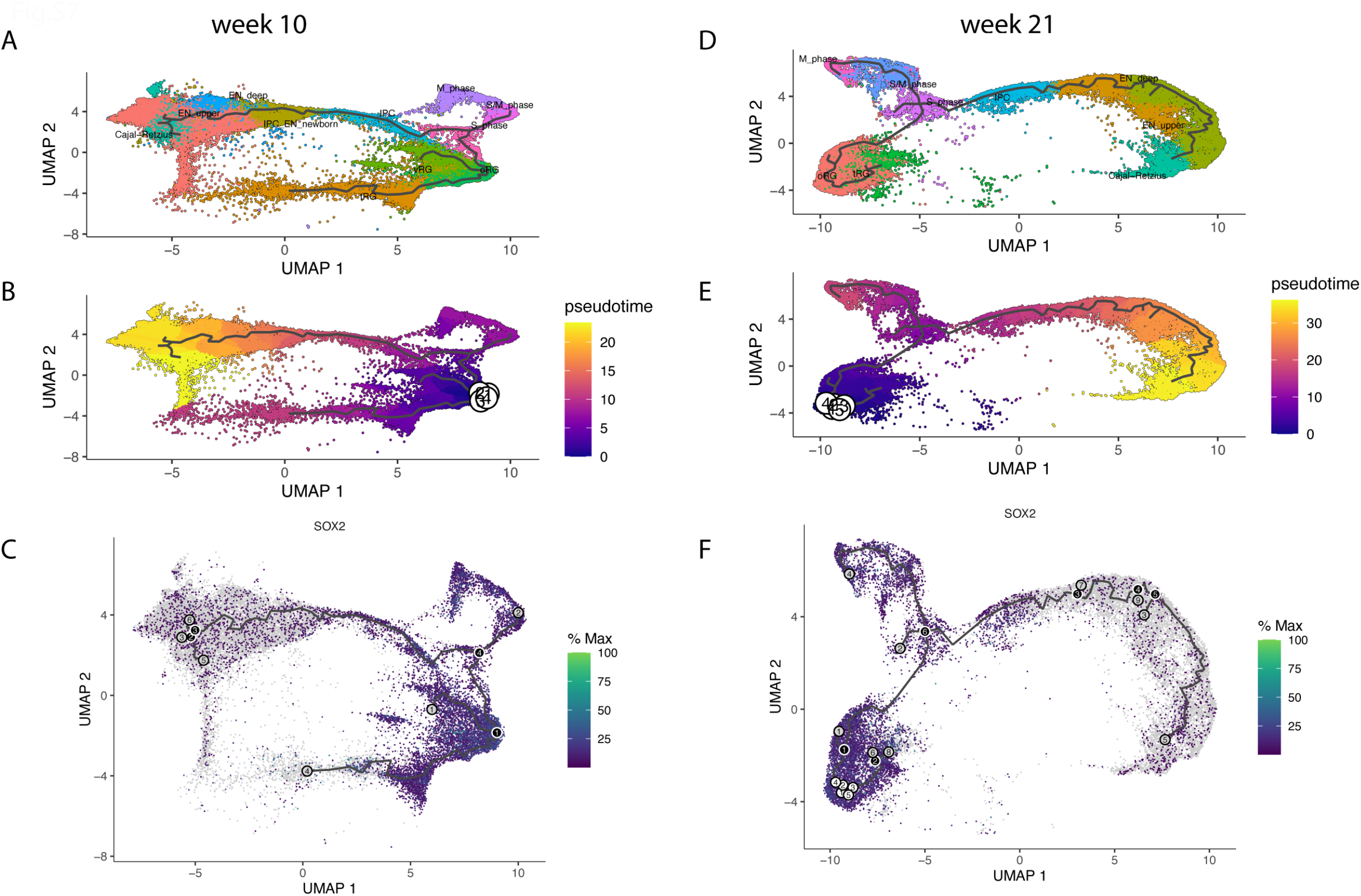
*PTEN* WT/I135L mutation accelerates the maturation of the upper layer neurons in the ASD genetic background, related to Figure 6. (A) UMAP plot of the cell types used for Monocle3 pseudotime analysis for week 10 organoids. (B) Pseudotime trajectory on UMAP plot for week 10 organoids. SOX2^+^ radial glia cells were used as root cells. (C) Relative expression of SOX2 on the UMAP plot for week 10 dataset. (D) UMAP plot of the cell types used for Monocle3 pseudotime analysis for week 21 dataset. (E) Pseudotime trajectory on UMAP plot for week 21 dataset. (F) Relative expression of SOX2 on the UMAP plot for week 21 dataset.

